# Identification of non-canonical peptides with moPepGen

**DOI:** 10.1101/2024.03.28.587261

**Authors:** Chenghao Zhu, Lydia Y. Liu, Annie Ha, Takafumi N. Yamaguchi, Helen Zhu, Rupert Hugh-White, Julie Livingstone, Yash Patel, Thomas Kislinger, Paul C. Boutros

## Abstract

Proteogenomics is limited by challenges of modeling the complexities of gene expression. We create moPepGen, a graph-based algorithm that comprehensively generates non-canonical peptides in linear time. moPepGen works with multiple technologies, in multiple species and on all types of genetic and transcriptomic data. In human cancer proteomes, it enumerates previously unobservable noncanonical peptides arising from germline and somatic genomic variants, noncoding open reading frames, RNA fusions and RNA circularization.

## Main Text

A single stretch of DNA can give rise to multiple protein products through genetic variation and through transcriptional, post-transcriptional and post-translational processes, such as RNA editing, alternative splicing and RNA circularization^1–4^. The number of potential proteoforms rises combinatorically with the number of possibilities at each level, so despite advances in proteomics technologies^5,6^, much of the proteome is undetected in high-throughput studies^7^.

The most common strategies to detect peptide sequences absent from canonical reference databases^7–9^ (*i.e.*, non-canonical peptides; **Supplementary Note 1**), are *de novo* sequencing and open search. Despite continued algorithmic improvements, these strategies are computationally expensive, have elevated false-negative rates, and lead to difficult data interpretation and variant identification issues^10,11^. As a result, the vast majority of proteogenomic studies use non-canonical peptide databases that have incorporated DNA and RNA alterations^7^. These databases are often generated using DNA and RNA sequencing of the same sample, and this improves error rates relative to community-based databases (*e.g.*, UniProt^12^, neXtProt^13^ and the Protein Mutant Database^14^) by focusing the search space^7,15^.

This type of sample-specific proteogenomics relies on the ability to predict all potential protein products generated by the complexity of gene expression. Modeling transcription, translation and peptide cleavage to fully enumerate the combinatorial diversity of non-canonical peptides is computationally demanding. To simplify the search-space, existing methods have focused on generating peptides caused by individual variants or variant types^16–33^, greatly increasing false negative rates and even potentially resulting in false positive detections if the correct peptide is absent from the database (**Extended Data Table 1**). To fill this gap, we created a graph-based algorithm for the exhaustive elucidation of protein sequence variations and subsequent *in silico* non-canonical peptide generation. This method is moPepGen (<u>m</u>ulti-<u>o</u>mics <u>Pep</u>tide <u>Gen</u>erator; **Figure 1a**).

**Figure 1:**
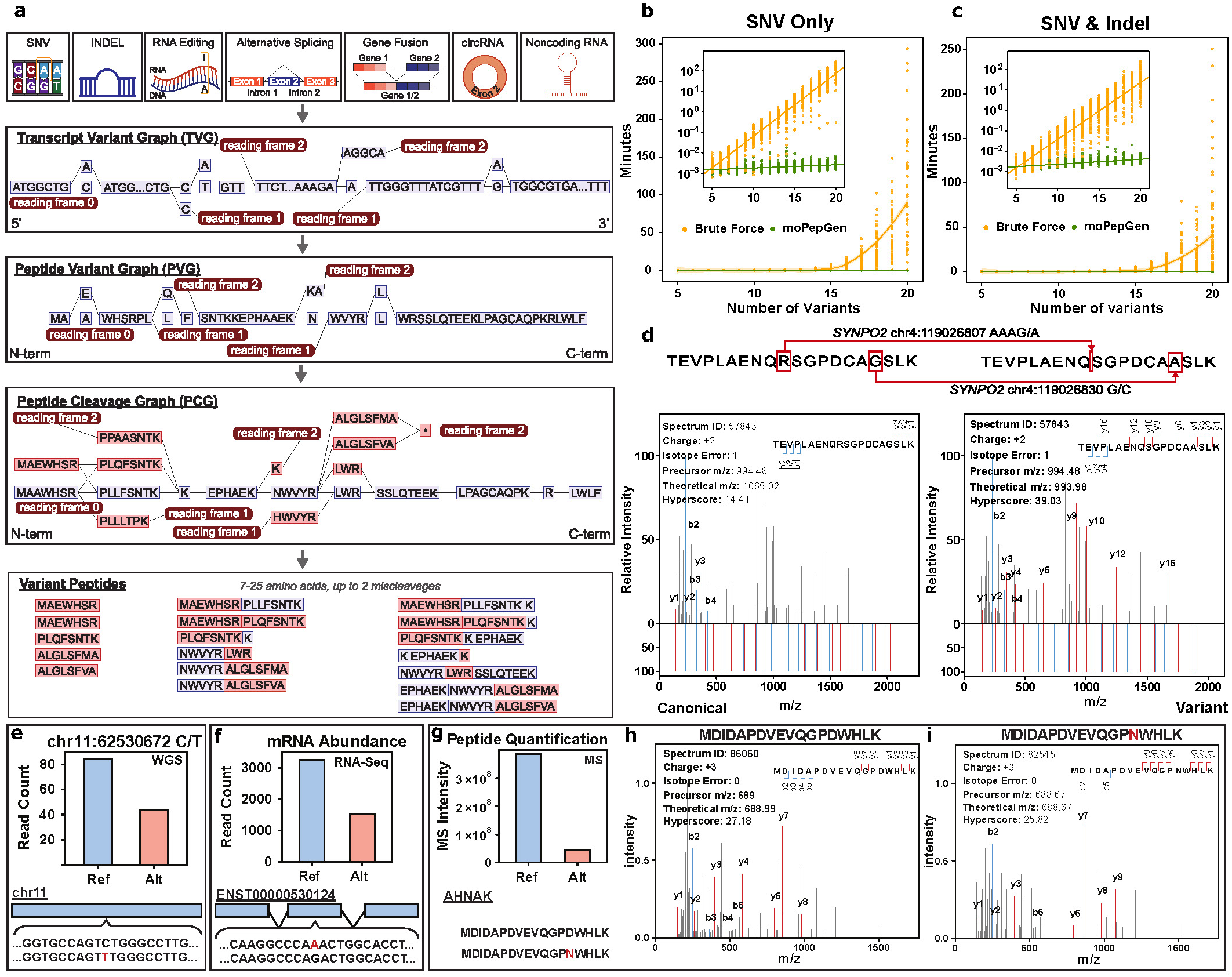
moPepGen is a graph-based algorithm that uncovers non-canonical peptides with variant combinations. **a**) moPepGen algorithm schematic. moPepGen is a graph-based algorithm that generates databases of non-canonical peptides that harbour genomic and transcriptomic variants (*e.g.,* single nucleotide variant (SNV), small insertion and deletion (INDEL), RNA editing, alternative splicing, gene fusion and circular RNA (circRNA)) from coding transcripts, as well as from novel open reading frames of noncoding transcripts. **b**) and **c**) moPepGen achieves linear runtime complexity when fuzz testing with SNVs only (**b**) and with SNVs and indels (**c**), based on 1,000 simulated test cases in each panel. **d**) A variant peptide from *SYNPO2* that harbours a small deletion and an SNV. Fragment ion mass spectrum from peptide-spectrum match (PSM) of the non-canonical peptide harbouring two variants (top, both) is compared against the canonical peptide theoretical spectra (left, theoretical spectra at the bottom) and against the variant peptide theoretical spectra (right, bottom). Fragment ion matches are colored, with b-ions in blue and y-ions in red. **e-g**) A somatic SNV D1249N in *AHNAK* was detected in DNA sequencing of a prostate tumour (CPCG0183) at chr11:62530672 (**e**), in RNA-sequencing (**f**) and as the non-canonical peptide MDIDAPDVEVQGP**N**WHLK (**g**). **h-i**): Fragment ion mass spectrum from PSM of the canonical peptide MDIDAPDVEVQGP**D**WHLK (**h**) and the non-canonical peptide (**i**). *m/z*: mass-to-charge ratio.

moPepGen captures peptides that harbour any combination of small variants (*e.g.*, single nucleotide polymorphisms (SNPs), small insertions and deletions (indels), RNA editing sites) occurring on canonical coding transcripts, as well as on non-canonical transcript backbones resulting from novel open reading frames (ORFs), transcript fusion, alternative splicing and RNA circularization (**Supplementary Figure 1**). It performs variant integration, *in silico* translation and peptide cleavage in a series of three graphs for every transcript, enabling systematic traversal across every variant combination (**Online Methods**; **Extended Data Figure 1a-d**). All three reading frames are explicitly modeled for both canonical coding transcripts and non-canonical transcript backbones to efficiently capture frameshift variants and facilitate three-frame ORF search (**Extended Data Figure 2a**). Alternative splicing events (*e.g.*, retained introns, *etc.*) and transcript fusions are modeled as subgraphs with additional small variants (**Extended Data Figure 2b**). Graphs are replicated four times to fully cover peptides of back-splicing junction read-through in circular RNAs (circRNAs; **Extended Data Figure 2c-d**). moPepGen outputs non-canonical peptides that cannot be produced by the chosen canonical proteome database. It documents all possible sources of each peptide to eliminate redundancy - for example where different combinations of genomic and transcriptomic events can produce the same non-canonical peptide.

We first validated moPepGen using 1,000,000 iterations of fuzz testing (**Supplementary Figure 2**). For each iteration, a transcript model, its nucleotide sequence, and a set of variants composed of all supported variant types were simulated. Then non-canonical peptides generated by moPepGen were compared to those from a ground-truth brute-force algorithm. moPepGen demonstrated perfect accuracy and linear runtime complexity (4.7 × 10^-3^ seconds per variant) compared to exponential runtime complexity for the brute-force method (**Figure 1b-c**). A comprehensive non-canonical peptide database of human germline polymorphisms was generated with 15 GB of memory in 3.2 hours on a 16-core compute node; the brute-force method was unable to complete this task.

Having established the accuracy of moPepGen, we next compared it to two popular custom database generators, customProDBJ^18^ and pyQUILTS^22^. We tested all three methods on five prostate tumours with extensive multi-omics characterization^34–36^. We first evaluated the simple case of germline and somatic point mutations and indels. Most peptides (84.0 ± 0.9% (median ± MAD (median absolute deviation))) were predicted by all three methods, with moPepGen being modestly more sensitive (**Extended Data Figure 3a**). Next, we considered the biological complexity of alternative splicing, RNA editing, RNA circularization and transcript fusion. Only moPepGen was able to evaluate peptides generated by all four of these processes, and therefore 80.2 ± 2.1% (median ± MAD) of peptides were uniquely predicted by moPepGen (**Extended Data Figure 3b**). By contrast only 3.2% of peptides were not predicted by moPepGen, and these corresponded to specific assumptions around the biology of transcription and translation made by other methods (**Extended Data Figure 3c**; **Online Methods**). By generating a more comprehensive database, moPepGen enabled the unique detection of 53.7 ± 12.2% (median ± MAD) more peptide hits from matched proteomic data (**Extended Data Figure 3d**). An example of a complex variant peptide identified only by moPepGen is the combination of a germline in-frame deletion followed by a substitution in *SYNPO2* (**Figure 1d**). In addition, moPepGen’s clear variant annotation system readily enables peptide verification across the central dogma. For example, the somatic mutation D1249N in *AHNAK* was detected in ∼30% of both DNA and RNA reads and was detected by mass spectrometry (MS; **Figure 1e-i**), confirmed by three search engines. Taken together, these benchmarking results demonstrate the robust and comprehensive nature of moPepGen.

To illustrate the use of moPepGen for proteogenomic studies, we first evaluated it across multiple proteases (**Extended Data Figure 4a**). Using independent conservative control of false discovery rate (FDR) across canonical and custom databases (**Online Methods**; **Supplementary Figure 3**)^7,36^, we focused on detection of novel ORFs (*i.e.*, polypeptides from transcripts canonically annotated as noncoding) across seven proteases^37^ in a deeply fractionated human tonsil sample^38^ (**Supplementary Table 1**). moPepGen enabled the detection of peptides from 1,787 distinct ORFs previously thought to be noncoding, and these peptides were most easily detected with the Arg-C protease (**Extended Data Figure 4b**), suggesting alternative proteases may enhance noncoding ORF detection (**Extended Data Figure 4c**). In total, 184 noncoding ORFs were detected across four or more proteomic preparation methods in this single sample, demonstrating that moPepGen can reliably identify novel proteins (**Extended Data Figure 4d-e**).

We next sought to demonstrate that moPepGen can benefit analyses in different species by studying germline variation in the C57BL/6N mouse^39,40^. Using strain-specific germline SNPs and indels from the Mouse Genome Project^39,40^, moPepGen predicted 5,481 non-canonical peptides arising from variants in protein-coding genes and 15,475 peptides from noncoding transcript novel ORFs (**Extended Data Figure 5a**). Across the proteomes of three bulk tissues (cerebellum, liver and uterus), we detected 18 non-canonical peptides in protein-coding genes and 343 from noncoding ORFs (**Extended Data Figure 5b-d**; **Supplementary Table 2**). Thus, moPepGen can support proteogenomics in non-human studies to identify variants of protein-coding genes and novel proteins.

To evaluate the use of moPepGen for somatic variation, we analyzed 375 human cancer cell line proteomes with matched somatic mutations and transcript fusions^41,42^ (**Supplementary Data**). moPepGen processed each cell line in 2:58 minutes (median ± 1:20 minutes, MAD), generating 2,683 ± 2,513 (median ± MAD) potential non-canonical variant peptides per cell line. The number of predicted variant peptides varied strongly with tissue of origin, ranging from median of 838 to 16,255 (**Figure 2a**) and was driven largely by somatic mutations in protein-coding genes and by fusion events in noncoding genes (**Extended Data Figure 6a-c**). Searching the cell line proteomes identified 39 ± 27 (median ± MAD) non-canonical peptides per cell line (**Online Methods**; **Supplementary Figure 4**). The majority of these were derived from noncoding transcript ORFs (**Extended Data Figure 6d**; **Supplementary Table 3**). Variant peptides from coding somatic mutations were more easily detected than those from transcript fusion events (**Extended Data Figure 6e-f**). 26 genes had variant peptides detected in cell lines from three or more tissues of origin, including the cancer driver genes *TP53, KRAS* and *HRAS* (**Figure 2b**). Peptide evidence was also found for fusion transcripts involving cancer driver genes like *MET* and *STK11* (**Extended Data Figure 6g-h**). We validated non-canonical peptide-spectrum matches (PSMs) by predicting tandem mass (MS2) spectra using Prosit^43^ and verifying that variant peptide MS2 spectra correlated better with predictions based on the matched non-canonical peptide sequences than predictions based on their canonical peptide counterparts (**Online Methods**; **Extended Data Figure 6i**). Coding variant peptide PSMs also showed high cross-correlations with their Prosit-predicted variant MS2 spectra, on par with those of canonical PSMs and their canonical spectra (**Extended Data Figure 6j**). Thus, moPepGen can effectively and rapidly detect variant peptides arising from somatic variation. These variant peptides may also prove to harbour functional consequences in future studies. Genes, such as *KRAS*, trended towards greater essentiality for cell growth in multiple cell lines with non-canonical peptide hits, and the effects may be independent of gene dosage (**Extended Data Figure 7a-c**). Across cell lines, detected variant peptides were also predicted to give rise to 416 putative neoantigens (3.0 ± 1.5, median ± MAD per cell line; **Extended Data Figure 7d**; **Supplementary Table 4**), including recurrent neoantigens in *KRAS*, *TP53* and *FUPBP3* (**Extended Data Figure 7e**).

**Figure 2:**
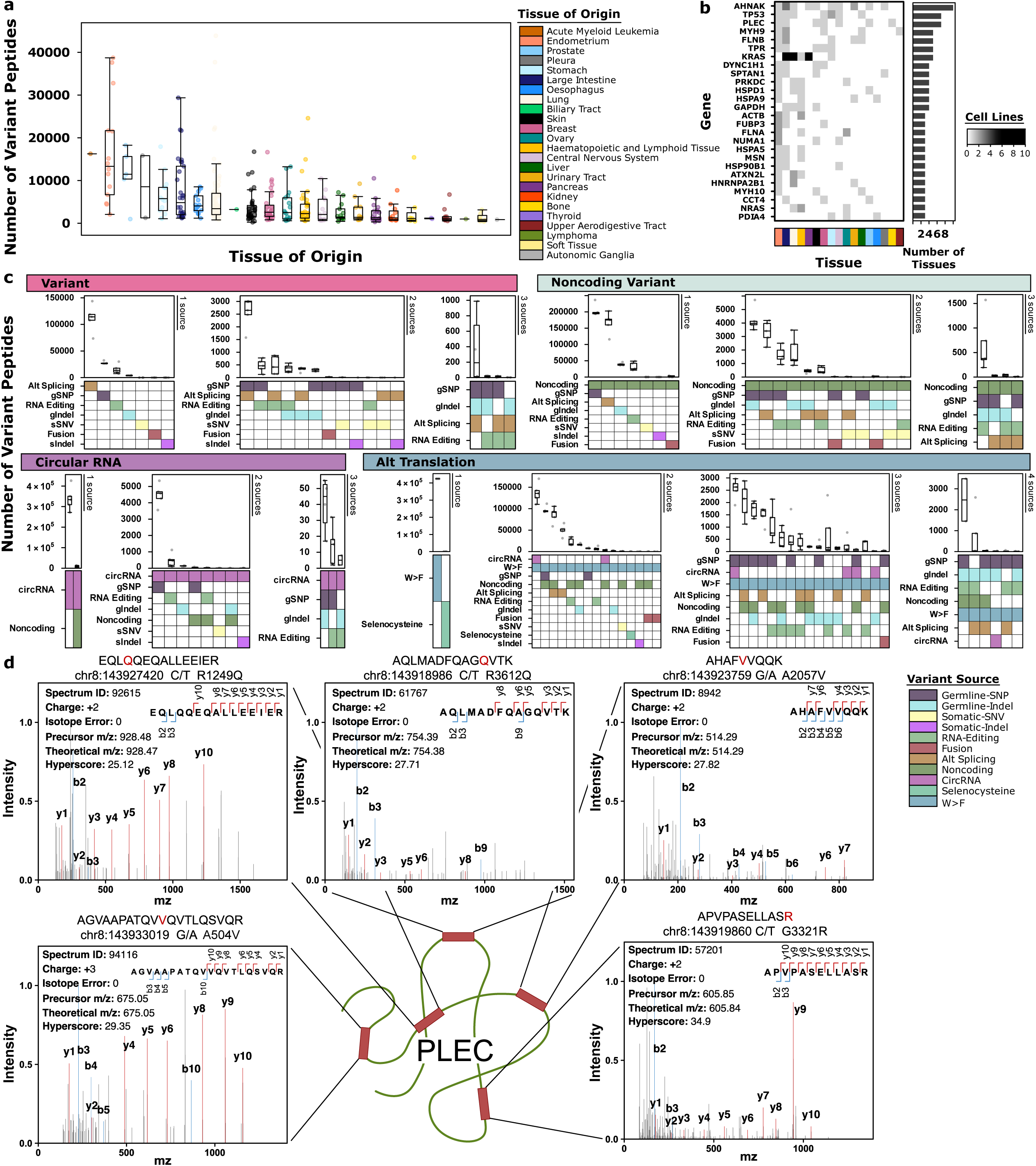
moPepGen generates comprehensive non-canonical databases that support proteogenomic analysis. **a)** Sizes of variant peptide databases generated by moPepGen using somatic single nucleotide variants, small insertions and deletions and transcript fusions for 376 cell lines from the Cancer Cell Line Encyclopedia project. Color indicates cell line tissue of origin. The number of cell lines per tissue of origin is provided in **Supplementary Table 8**. **b**) Genes with variant peptides detected in cell lines across three or more tissues of origin (bottom covariate). The barplot shows number of recurrences across tissues and color of heatmap indicates number of cell lines. **c**) Number of non-canonical peptides from different variant combinations (bottom heatmap) generated using genomic and transcriptomic data from five primary prostate tumours (n = 5), shown across four tiers of custom databases and grouped by the number of variant sources in combination. Alternative translation (Alt Translation) sources with ≥ 10 peptides are visualized. gSNP: germline single nucleotide polymorphism; gIndel: germline small insertion and deletion (indel); sSNV: somatic single nucleotide variant; sIndel: somatic indel; circRNA: circular RNA; W>F: tryptophan-to-phenylalanine. **d**) Five variant peptides detected in one prostate tumour (CPCG0183) from the protein plectin (PLEC). Fragment ion matches are colored, with b-ions in blue and y-ions in red. *m/z*: mass-to-charge ratio. All boxplots show the first quartile, median, to the third quartile, with whiskers extending to furthest points within 1.5× the interquartile range.

We next sought to demonstrate the use of moPepGen in data-independent acquisition (DIA) MS using eight clear cell renal cell carcinoma tumours with matched whole-exome sequencing, RNA-sequencing and DIA proteomics^44^. In each tumour, moPepGen predicted 157,016 ± 34,215 (median ± MAD) unique variant peptides from protein-coding genes (**Extended Data Figure 8a**). Using a Prosit-generated spectral library, we detected 307 ± 112 (median ± MAD) variant peptides in each tumour using DIA-NN^45^ (**Extended Data Figure 8b**; **Supplementary Table 5**). Germline-SNP and alternative splicing were the most common sources of detected variant peptides (**Extended Data Figure 8c-d**). Non-canonical peptides derived from RNA editing events were detected in 21 genes (**Extended Data Figure 8e-i**). Thus, moPepGen can enable the detection of variant peptides from DIA proteomics.

Finally, to demonstrate the use of moPepGen on complex and comprehensive gene expression data, we analyzed five primary prostate cancer samples with matched DNA whole-genome sequencing, ultra-deep ribosomal-RNA-depleted RNA-sequencing and MS-based proteomics^34–36^. moPepGen generated 1,382,666 ± 64,281 (median ± MAD) unique variant peptides per sample, spanning 115 variant combination categories (**Figure 2c**). Searching this database resulted in the detection of 206 ± 56 (median ± MAD) non-canonical peptides per sample, with 138 ± 28 (median ± MAD) derived from protein-coding genes (**Extended Data Figure 9a**, **Supplementary Table 6**). The distribution of intensities and Comet expectation scores of non-canonical PSMs closely resembled that of canonical PSMs and was distinct from all decoy hits (**Supplementary Figure 5**), lending confidence in our non-canonical peptide detection. All samples harboured proteins containing multiple variant peptides (9 ± 1.5, median ± MAD proteins per tumour; range 2-6 variant peptides per protein; **Figure 2d**). Some detected peptides harboured multiple variants, including two from prostate-specific antigen (PSA from the *KLK3* gene; **Extended Data Figure 9b**). Germline SNPs were the major common cause of variant peptides on coding transcripts and alternative splicing events were the most common cause on noncoding transcripts (**Extended Data Figure 9c-e**). Nine genes showed recurrent detection of peptides caused by circRNA back-splicing (**Extended Data Figure 9f-g**), with 36/78 circRNA PSMs validated by *de novo* sequencing (**Supplementary Table 7**)^46^. These recurrent circRNA-derived peptides were verified in five additional prostate tumours (**Supplementary Figure 6**). We also detected four peptides from noncoding transcripts with the recently reported tryptophan-to-phenylalanine substitutants^47^. Thus, moPepGen can identify peptides resulting from highly complex layers of gene expression regulation.

moPepGen is a computationally efficient algorithm that enumerates transcriptome and proteome diversity across arbitrary variant types. It enables the detection of variant and novel ORF peptides across species, proteases and technologies. moPepGen integrates into existing proteomic analysis workflows, and can broadly enhance proteogenomic analyses for many applications.

## Online Methods

### Transcript Variant Graph

A transcript variant graph (TVG) is instantiated for each transcript, incorporating all associated variants. In a TVG, nodes are transcript fragments with reference or alternative nucleotide sequences, while edges are the opening or closing of variant nodes connecting them to the reference sequence, or the elongation of reference sequences. The TVG starts with three linear nodes of the entire transcript sequence representing the three reading frames, offset by 0, 1, or 2 nucleotides from the transcript 5′ end. A variant is incorporated into the graph by breaking the node at the variant’s start and end positions and attaching a new node with the alternative sequence to the new upstream and downstream nodes. An in-frame variant is represented as a node with incoming and outgoing nodes in the same reading frame subgraph, while frameshifting variants have incoming nodes and outgoing nodes in different reading frames. The outgoing reading frame index equals to (*S*_*ref*_ − *S*_*alt*_) *mod* 3, where *S*_*ref*_ is the length of the reference sequence and *S*_*alt*_ is the length of the alternative sequence. For transcripts with an annotated known canonical open reading frame (ORF), variants are only incorporated into the subgraph of the appropriate reading frame (**Extended Data Figure 2a**). If frameshifting variants are present, downstream variants are also incorporated into the subgraphs of the outgoing frameshift nodes. For transcripts without an annotated ORF, all variants are incorporated into all three reading frames (**Extended Data Figure 2a**). Large insertions and substitutions as the result of alternative splicing events (*e.g.*, retained introns, alternative 3’/5’ splicing, *etc.*) are represented as subgraphs that can carry additional variants (**Extended Data Figure 2b)**.

### Variant Bubbles and Peptide Variant Graph

After the TVG has been populated with all variants, nodes that overlap with each other in transcriptional coordinates are aligned to create variant bubbles within which all nodes point to the same upstream and downstream nodes (**Extended Data Figure 1b**). This is done by first finding connection nodes in the TVG, the reference nodes without any variants that connect two variant bubbles after they are aligned. The root node is the first connection node, and the next connection node is found by looking for the first commonly connected downstream node with length of five or more nucleotides that is outbound to more than one node (**Supplementary Note 2**; **Supplementary Figure 7**). Nodes between the two connection nodes are then aligned to form a variant bubble by generating all combinations of merged nodes so that they all point to the same upstream and downstream nodes (**Extended Data Figure 1b**). Overlapping variants in the variant bubble are automatically eliminated because they are disjoint. The sequence lengths of nodes in the variant bubble are also adjusted by taking nucleotides from the commonly connected upstream and downstream nodes to ensure that they are multiples of three. A peptide variant graph (PVG) is then instantiated by translating the nucleotide sequence of each TVG node into amino acid sequences.

### Peptide Cleavage Graph

A PVG is converted into a peptide cleavage graph (PCG), where each edge represents an enzymatic cleavage site (**Extended Data Figure 1c**). For connection nodes, all enzymatic cleavage sites are first identified, and the node is cleaved at each cleavage site. Because enzymatic cleavage site motifs can span over multiple nodes (for example, the trypsin exception of not cutting given K/P but cutting given WK/P), connection nodes are also merged with each downstream and/or upstream node and cut at additional cleavage sites if found. To optimize run time, different *merge-and-cleave* operations are used depending on the number of incoming and outgoing nodes, and the number of cleavage sites in a node (**Supplementary Figure 8**). Hypermutated regions where variant bubbles contain many variants and/or the lack of cleavage sites in connection nodes can result in an exponential increase in the number of nodes in the aligned variant bubble. We use a *pop-and-collapse* strategy, such that when *merge-and-cleave* is applied to a connection node, *x* number of amino acids are popped from the end of each node in the variant bubble. The popped nodes are collapsed if they share the same sequence. The *pop-and-collapse* operation is only applied when the number of nodes in a variant bubble exceeds a user-defined cutoff.

### Calling Variant Peptides

Variant peptides with the permitted number of miscleavages are called by traversing through the peptide cleavage graph. We use a *stage-and-call* approach that first visits all incoming nodes to determine the valid ORFs of a peptide node (**Supplementary Note 3**). *Stage-and-call* also allows cleavage-gain mutations and upstream frameshift mutations to be carried over to the downstream peptide nodes. Peptide nodes are then extended by merging with downstream nodes to call variant peptides with miscleavages (**Supplementary Note 4**). For noncoding transcripts, novel ORF start sites, including those caused by start-gain mutations, are found by looking for any methionine (M) in all three subgraphs. Terminology used in subsequent sections, including canonical and non-canonical database, variant peptides, non-canonical peptides, and proteoform, is defined in **Supplementary Note 1**.

### Fusion and Circular Transcripts

Most fusion transcript callers detect fusion events between genes, causing ambiguity of which transcripts of the genes are involved in a particular fusion event. We took the most comprehensive approach and endeavored to capture all possible variant peptides by assuming that a fusion event could happen between any transcript of the donor and accepter genes. Fusion transcripts are considered as novel backbones in graph instantiation, with an individual graph instantiated for each donor and acceptor transcript pair. Single nucleotide variants (SNVs) and small insertion/deletions (indels) of both donor and acceptor transcripts are incorporated into the TVG. The translated and cleaved PCG is then traversed to call variant peptides, identical to a canonical transcript backbone. If the fusion breakpoint occurs in an intron, the intronic nucleotide sequence leading up to or following the breakpoint is retained as unspliced, and its associated intronic variants are included. The ORF start site of the donor transcript is used if exists when calling variant peptides. The fusion transcript is treated as a noncoding transcript if the donor transcript is annotated as noncoding.

Similar to fusion transcripts, circular RNAs (circRNAs) are treated as novel backbones, with an individual graph instantiated for each circRNA (Extended Data Figure 2c). A circular variant graph (CVG, a counterpart to TVG) is instantiated by connecting the linear sequence of the circRNA onto itself at the back-splice junction and incorporating SNVs and indels. Novel peptides can theoretically be translated from circRNAs if a start codon is present, by ribosome readthrough across the back-splicing junction site. If the circRNA length is not a multiple of three nucleotides, translation across the back-splicing site induces a frameshift. Without a stop codon, the ribosome may traverse the circRNA up to three times before the amino acid sequence repeats. Therefore, moPepGen extends the circular graph linearly by appending three copies of each reading frame as a subgraph to account for frameshifts. The extended graph is then translated to a PVG and converted to a PCG. Variant peptides are called by treating every circRNA as a noncoding transcript and scanning all novel start codons in all three reading frames.

### Biological Assumptions for Edge Cases

moPepGen applies various assumptions to selectively include or exclude certain variant events or peptides (**Extended Data Figure 3c**). Start-codon-altering variants are excluded due to the uncertainty around whether and where translation will still occur. Similarly, splice-site-altering variants are omitted due to the complexity of splicing determinants, which can result in skipping to the next canonical or non-canonical splice site. We terminate translation at the last complete peptide when stop codons are unknown, as incomplete transcript annotations create ambiguity in downstream sequences, obscuring enzymatic cleavage sites. Stop-codon-altering variants do not extend translation beyond the transcript, as the downstream genomic region is not assumed to be part of the RNA transcript.

### GVF File Format and Parsers

Genomic (single nucleotide polymorphisms (SNP), SNV, indel) and transcriptomic variants (fusion transcripts, RNA editing sites, alternative splicing transcripts, circRNAs) are first converted into gene-centric entries for each transcript that they impact. We defined a gene-based GVF (genetic variant format) derived from VCF (variant calling format) to store all relevant information for each variant, including the gene ID and offset. moPepGen includes built-in parsers to convert variant caller outputs into GVFs. SNPs, SNVs and indels require the annotation via Variant Effect Predictor (VEP) for compatibility with the parseVEP module. Parsers for fusion, alternative splicing, RNA editing and circRNA operate directly on native outputs. moPepGen is implemented in Python and supports easy extension and addition of new parsers. The full Nextflow pipeline (https://github.com/uclahs-cds/pipeline-call-NonCanonicalPeptide)^48^, automates data preprocessing, peptide prediction and database tiering, with optional transcript abundance filtering^49,50^. Our DNA data processing pipeline is described elsewhere^51^.

### Fuzz Testing and Brute Force Algorithm

To validate moPepGen, we implemented a fuzz testing framework where transcripts with varying properties (*e.g.*, coding status, strand, selenocysteine and start or stop codon position) and artificial sequences are simulated. Each is paired with simulated variants across all supported types. The resulting peptides are compared against those generated by a brute force algorithm, which iterates through all possible variant combinations to identify non-canonical peptides. The brute force algorithm also performs three-frame translation for noncoding transcripts. Fuzz testing and the brute force algorithm are included in the moPepGen package.

### Datasets

#### Cancer Cell Line Encyclopedia proteome

Proteomics characterization of 375 cell lines from the Cancer Cell Line Encyclopedia (CCLE) was obtained from Nusinow *et al.*, 2020^52^. Fractionated raw mass spectrometry (MS) data were downloaded from MassIVE (project ID: MSV000085836). Somatic SNVs and indels, and fusion transcript calls were downloaded from the DepMap portal (https://depmap.org/portal, 22Q1). Somatic SNVs and indels were converted to GRCh38 coordinates from hg19 using CrossMap (v0.5.2)^53^. Gene and transcript IDs were assigned to each SNV/indel using VEP (v104) ^54^ with genomic annotation GTF downloaded from GENCODE (v34)^55^. Fusion results were aligned to the GENCODE v34 reference by first lifting over the fusion coordinates to GRCh38 using CrossMap (v0.5.2). After lift-over, the records were removed if the donor or acceptor breakpoint location was no longer associated with the gene, if either breakpoint dinucleotides did not match with the reference, or if either gene ID was not present in GENCODE (v34).

#### Mouse proteome

MS-based proteome of mouse strain C57BL/6N was obtained from Giansanti *et al.*, 2022^40^. Fractionated raw MS data of the liver, uterus and cerebellum proteomes were downloaded from the PRIDE repository (project ID: PXD030983). Germline SNPs and indels were obtained from the Mouse Genomes Project^39^ with GRCm38 VCFs downloaded from the European Variation Archive (accession: PRJEB43298). Germline SNPs and indels were annotated using VEP (v102) against Ensembl GRCm38 (v102)^56^.

#### Alternative protease and fragmentation proteome

A human tonsil tissue processed using ten different combinations of proteases and peptide fragmentation methods (ArgC_HCD, AspN_HCD, Chymotrypsin_CID, Chymotrypsin_HCD, GluC_HCD, LysC_HCD, LysN_HCD, Trypsin_CID, Trypsin_ETD, Trypsin_HCD) was obtained from Wang *et al.*, 2019^38^. Fractionated raw mass spectrometry data were downloaded from the PRIDE repository (project ID: PXD010154).

#### DIA Proteome

Data-independent acquisition (DIA) proteomic data from eight clear cell renal cell carcinoma (ccRCC) samples were obtained from Li *et al.*, 2023^44^. Raw mass spectrometry data were retrieved from the Proteomic Data Commons (PDC, PDC000411). WXS and RNA-seq BAM files were obtained from Genomic Data Commons (GDC, Project: CPTAC-3, Primary Site: Kidney). WXS data was processed using a standardized pipeline to identify germline SNPs, somatic SNVs and indels^51^. BAM files were reverted to FASTQ using Picard toolkit (v2.27.4) and SAMtools (v1.15.1)^57^, re-aligned to GRCh38 using BWA-MEM2 (v2.2.1)^58^, and calibrated using BQSR and IndelRealignment from GATK (v4.2.4.1)^59^. Germline SNPs and indels were called following GATK (v4.2.4.1) best practices^59,60^, while somatic SNVs and indels were called using Mutect2 (from GATK v4.5.0.0), followed by annotation with VEP (v104)^61^ against GENCODE v34. RNA-seq BAM files were converted to FASTQ using Picard toolkit (v2.27.4) and SAMtools (v1.15.1) and re-aligned to GRCh38.p13 with GENCODE v34 GTF using STAR (2.7.10b)^62^. Transcript fusion events were called using STAR-Fusion (v1.9.1)^63^, alternative splicing events were called using rMATS (v4.1.1)^64^, and RNA editing sites were called using REDItools2 (v1.0.0)^65^ using paired RNA and DNA BAMs.

#### Prostate cancer proteome

The proteomics characterization of five prostate cancer tissues were obtained from Sinha *et al.*, 2019^36^. Raw mass spectrometry data were downloaded from MassIVE (project ID: MSV000081552). Germline SNPs and indels, as well as somatic SNVs and indels were obtained from the ICGC Data Portal (Project code: PRAD-CA). Variants were indexed using VCFtools (v0.1.16)^66^ and converted to GRCh38 using Picard toolkit (v2.19.0), followed by chromosome name mapping from the Ensembl to the GENCODE system using BCFtools (v1.9-1)^67^. Mutations were annotated using VEP (v104)^61^ against GENCODE (v34). Raw mRNA sequencing data were obtained from Gene Expression Omnibus (accession: GSE84043). Transcriptome alignment was performed using STAR (v2.7.2) to reference genome GRCh38.p13 with GENCODE (v34) GTF and junctions were identified by setting the parameter --chimSegmentMin 10^68^. CIRCexplorer2 (v.2.3.8) was used to parse and annotate junctions for circular RNA detection^69^. Fusion transcripts were called using STAR-Fusion (v1.9.1)^63^. RNA editing sites were called using REDItools2 using paired RNA and DNA BAMs (v1.0.0)^65^. Alternative splicing transcripts were called using rMATS (v4.1.1)^64^.

### Canonical Database Search

All MS raw files (.raw) were converted to mzML using ProteoWizard (3.0.21258)^70^. The GRCh38 human and the GRCm38 mouse canonical proteome databases were obtained from GENCODE (v34) and Ensembl (v102), respectively, with common contaminants^71^ added and reversed sequences appended for target-decoy false discovery rate (FDR) control. Database searches were performed using Comet (v2019.01r5)^72^ with static modifications of cysteine carbamidomethylation, and up to three variable modifications (methionine oxidation, protein N-terminus acetylation, peptide N-terminus pyroglutamate formation), under full trypsin digestion with up to two miscleavages (except for the tonsil samples processed with alternative enzymes), for peptide lengths 7–35. For CCLE, static modification of tandem mass tag (TMT; 10plex) on the peptide N-terminus and lysine residues and variable modification of TMT on serine residues were additionally included, following the original study. CCLE data were searched in low resolution with 20 ppm precursor mass tolerance, 0.5025 Da fragment mass tolerance, and clear TMT *m/z* range, following the original publication. All other datasets used high-resolution label-free quantification (LFQ), with precursor mass tolerance of 20 ppm (mouse), 10 ppm (tonsil) and 30 ppm (prostate), and fragment mass tolerance of 0.025 Da for tonsil or 0.01 Da otherwise, following original publications. Tonsil proteomes were searched with the appropriate protease used in sample preparation, with a maximum of two miscleavages for Lys-C and Arg-C, three miscleavages for Glu-C and Asp-N and four miscleavages for chymotrypsin, as in the original publication^38^. The eight DIA ccRCC proteomes were not searched against a canonical database.

Peptide level target-decoy FDR calculation was performed using the FalseDiscoveryRate module from OpenMS (v3.0.0-1f903c0)^73^ using the formula (D+1)/(T+D), where D and T are the numbers of decoy and target peptide-spectrum matches (PSMs), respectively. Peptides were filtered at 1% FDR, and PSMs were removed from the corresponding mzML for subsequent non-canonical database search. *Post-hoc* cohort-level FDR was calculated to verify an FDR cutoff smaller than 1%. Peptide quantification was performed using OpenMS FeatureFinderIdentification (v3.0.0-1f903c0)^74^ with “internal IDs only” and adjusted precursor mass tolerances as above, and otherwise default parameters. TMT quantification was performed using OpenMS IsobaricAnalyzer (v3.0.0-1f903c0), without isotope correction due to absence of correction matrix.

### Non-canonical Database Generation

Human (GRCh38) and mouse (GRCm38) reference proteomes were obtained from GENCODE (v34) and Ensembl (v102), respectively. Non-canonical peptide databases were generated with trypsin digestion of up to two miscleavages and peptide lengths 7-25, except for alternative protease samples. Alternative translation peptides were generated using *callAltTranslation*, including those with selenocysteine termination^75^ or W>F substitutants^47^. Peptides from noncoding ORFs were generated using *callNovelORF* with ORF order as min and with or without alternative translation. Noncoding ORF peptide databases were also generated for each alternative protease used in processing of the tonsil proteome, with appropriate number of maximum miscleavages as outlined above.

Non-canonical peptide databases were generated for 376 CCLE cell lines, 375 of which have non-reference channel proteomics characterization. This included all 10 cell lines in the bridge line and 366 non-reference cell lines with mutation data. Of the 378 non-reference channels across 42 plexes, three cell lines were duplicated, seven were in the bridge line, two didn’t have mutation or fusion information and additional eight didn’t have fusion information. Variant databases from all cell lines in a TMT plex, including the ten cell lines in the reference channel, were merged along with noncoding ORF peptides to generate plex-level databases. Plex-level databases were split into three tiers: “Coding” (SNVs, indels and fusion in coding transcripts), “Noncoding” (novel ORFs) and “Noncoding Variant” (SNVs, indels and fusion in noncoding transcripts).

Non-canonical peptide databases for the proteome of mouse strain C57BL/6N were generated by calling variant peptides based on germline SNPs and indels, followed by merging with the noncoding ORF peptides. The resulting non-canonical peptides were then split into “Germline” (variants in coding transcripts), “Noncoding” (novel ORFs) and “Noncoding-Germline” (variants in noncoding transcripts).

For the eight ccRCC tumours, sample-specific variant peptides were called from germline/somatic SNVs and indels, RNA editing, transcript fusion and alternative splicing. Resulting peptides were merged with noncoding ORF and alternative translation peptides and split into four tiers: “Variant” (variants in coding transcripts), “Noncoding” (novel ORFs), “Noncoding Variant” (variants in noncoding transcripts) and “Alt Translation” (selenocysteine termination and W>F substitutants^47^).

For the five prostate tumours, variant peptides were called from all available genomic and transcriptomic variants, including germline/somatic SNVs and indels, RNA editing sites, transcript fusions, alternative splicing and circRNA. These peptides were then merged with the noncoding ORF and alternative translation peptides and split into five tiers: “Variant” (variants in coding transcripts), “Noncoding” (novel ORFs), “Noncoding Variant” (variants in noncoding transcripts), “Circular RNA” (circRNA ORFs) and “Alt Translation” (selenocysteine termination and W>F substitutants^47^).

### Non-canonical Database Search

Non-canonical database searches were performed similarly to canonical proteome searches for each dataset, as described in detail above. Custom databases of peptide sequences were concatenated with the reverse sequence for FDR control. Non-canonical peptide searches with Comet (v2019.01r5) were set to “no cleavage” and did not permit protein N-terminus modifications or clipping of N-terminus methionine. Peptide-level FDR was set to 1% independently for each tier of non-canonical database, and PSMs of peptides that passed FDR were removed from the mzML for subsequent searches. *Post-hoc* cohort-level FDR was calculated to verify an FDR cutoff smaller than 1%. Each database tier thus had independent FDR control using database-specific decoy peptides, and a spectrum is excluded from subsequent searches after finding its most probable match. This strategy minimizes false-positives caused by joint FDR calculation with canonical peptides and enables a conservative detection of non-canonical peptides^7,76^. For CCLE, peptides were only considered for detection and quantitation for a cell line if they existed in the sample-specific database. For prostate tumours, additional searches were conducted with the same non-canonical databases using MSFragger (v3.3)^77^ and X!Tandem (v2015.12.15)^78^ with equivalent parameters for verification. For all datasets, quantified peptides were distinguished by charge and variable modifications, and detected but not quantified peptides were excluded from subsequent analysis.

### DIA Non-canonical Spectral Library Search

Raw files were converted to .mzML files using ProteoWizard (3.0.21258)^70^. Sample-specific variant peptide FASTA databases were generated using the aforementioned non-canonical database generation pipeline, with individual spectral libraries .msp files generated by Prosit ^79^. Prosit was configured with instrument type of LUMAS, collision energy of 34, and fragmentation method of HCD, with all default parameters otherwise. Searches were conducted using DIA-NN (v1.8.1)^45^ against the sample-specific predicted variant peptide spectral libraries with protein inference disabled, a q-value cutoff of 0.01, and “high precision” quantification.

### Neoantigen Prediction

Neoantigens were predicted from non-canonical peptides detected in CCLE proteomes. Cell line-specific *HLA* genotype was inferred using OptiType (v1.3.5)^80^ from WGS or WXS data. Detected non-canonical peptides from the “Coding” tier were converted to FASTA and analyzed using MHCflurry (v2.0.6)^81^ with default parameters and cell line-specific *HLA* genotypes.

### Statistical Analysis and Data Visualization

All statistical analysis and data visualization were performed in the R statistical environment (v4.0.3), with visualization using BoutrosLab.plotting.general (v6.0.2)^82^. All boxplots, except for **Extended Data Figure 7a**, show all data points, the median (center line), upper and lower quartiles (box limits), and whiskers extend to the minimum and maximum values within 1.5 times the interquartile range. In **Extended Data Figure 7a**, data are summarized as boxplots to improve visual clarity without individual points due to the large number of gene and cell line combinations. All comparisons were performed on biological replicates, defined as independent patients, tumours, or cell lines as appropriate to each analysis. Schematics were created in Inkscape (v1.0) and Adobe Illustrator (27.8.1), and figures were assembled using Inkscape (v1.0).

#### Gene Dependency Association Analysis

Gene dependency data from CCLE CRISPR screens were downloaded from the DepMap data portal (https://depmap.org/portal, 24Q2). Twelve cell lines with non-canonical peptide detections in proteomic data from at least ten genes were selected. The CERES scores^83^ of genes with non-canonical peptide hits were compared to those without, using the Mann-Whitney U-test. Additionally, pooled CERES scores across all genes and cell lines were compared between the two groups using the same test. For *KRAS*, CERES scores and RNA abundance were compared between cell lines with non-canonical peptide detections in proteomic data and those with only canonical peptides, using the Mann-Whitney U-test.

#### Spectrum Visualization and Validation

Target PSM experimental spectra were extracted from mzML files using pyOpenMS (v3.1.0)^84^ and visualized in R. Theoretical spectra were generated from the target peptide sequences using the TheoreticalSpectrumGenerator module of OpenMS and compared to the experimental spectra using hyperscores via the HyperScore module with consistent parameters (*e.g.*, fragment mass tolerance). Fragment ion matching between the experimental and theoretical spectra was performed using a similar approach to IPSA^85^. Theoretical spectra with predicted fragment peak intensities were generated using Prosit via Oktoberfest (v0.6.2)^86^ with parameters (e.g., fragmentation method and energy) matching the original publication^43^ and compared using cross-correlation^87^ with settings matching Comet searches (*e.g.*, fragment_bin_offset). To assess the distribution of cross-correlation values for variant peptide PSMs, we randomly selected 1,000 canonical PSMs from each of the 42 TMT-plexes as control. circRNA peptide PSMs were validated using the Novor algorithm through app.novor.cloud, using parameters (*e.g.*, fragmentation method, MS2 analyzer, enzyme, precursor and fragment mass tolerance) consistent with database searches^46^.

#### Cohort Level FDR

A *post-hoc* approach was employed to estimate the FDR threshold at the cohort level for each database tier. Within each sample and database tier, we first identified the target hit with the highest FDR value under the 1% threshold, denoted as FDR^i^.

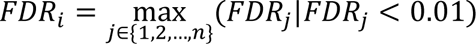

The number of decoy and target hits with FDR values less than FDR^i^ for each sample were tallied. The equivalent cohort-level FDR threshold was then calculated by dividing the total number of decoy hits by the total number of target and decoy hits across the cohort.

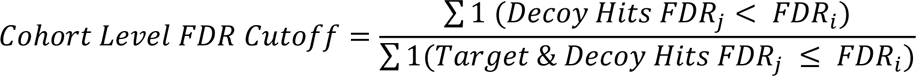

### Data Availability

Data supporting the conclusions of this paper is included within it and its supplementary files. The processed CCLE data are available at the DepMap portal (http://www.depmap.org). The raw WGS and WXS cell lines sequencing data are available at Sequence Read Archive (SRA) and European Genome-Phenome Archive (EGA) under access number PRJNA523380^41^ and EGAD00001001039^88^. The raw mass spectrometry proteomic data are publicly available without restrictions at the ProteomeXchange via the PRIDE partner repository under accession number PXD030304^42^ for cell lines, PXD030983^37^ for mouse strain C57BL/6N, and PXD010154^89^ for alternative protease and fragmentation analyses. The proteomic data for the five prostate tumour samples are freely available at UCSD’s MassIVE database under accession number MSV000081552^36^, whereas their raw WGS and RNA-seq data are available at EGA under accession EGAS00001000900^35^. Proteomic data for the eight ccRCC tumour samples are freely available at PDC under accession number PDC000411^44^, whereas the genomic and transcriptomic data are available at Genomic Data Commons (GDC, Project: CPTAC-3, Primary Site: Kidney) with dbGaP accession number phs001287, generated by the National Cancer Institute’s Clinical Proteomic Tumor Analysis Consortium (CPTAC).

## Acknowledgments

The authors thank members of the Boutros and Kislinger labs for their continued support, particularly Dr. Amanda Khoo, Meinusha Govindarajan and Dr. Matthew Waas. The authors also thank Dr. James Wohlschlegel from UCLA Proteome Research Center and Dr. Mathias Wilhelm from Technical University of Munich. This work was supported by the NIH *via* awards P30CA016042, U01CA214194, U2CCA271894, U24CA248265, P50CA092131 and R01CA244729, by the Canadian Cancer Society *via* an Impact Grant (705649) and by the Canadian Institute of Health Research *via* a Project Grant (PJT156357). CZ was supported by the UCLA Jonsson Comprehensive Cancer Center Fellowship Award. LYL was supported by a CIHR Vanier Fellowship and Ontario Graduate Scholarship. HZ was supported by a CIHR Doctoral Award. TK is supported through the Canadian Research Chair program. University Health Network was supported by the Ontario Ministry of Health and Long-Term care.

## Author Contributions

**Conceptualization:** C.Z., L.Y.L., T.K. and P.C.B.

**Software:** C.Z. and L.Y.L.

**Formal analysis:** C.Z., L.Y.L., A.H., T.N.Y., H.Z., R.H., J.L. and Y.P.

**Data Curation:** C.Z., L.Y.L., A.H., T.N.Y., H.Z., R.H., J.L. and Y.P.

**Visualization:** C.Z., L.Y.L. and A.H.

**Writing - Original Draft:** C.Z., L.Y.L., T.K. and P.C.B.

**Writing - Review & Editing:** All authors.

## Code Availability

moPepGen is publicly available at: https://github.com/uclahs-cds/package-moPepGen^90^. Data processing, analysis and visualization scripts are available upon request.

## Extended Data Figure Legends

**Extended Data Figure 1:**
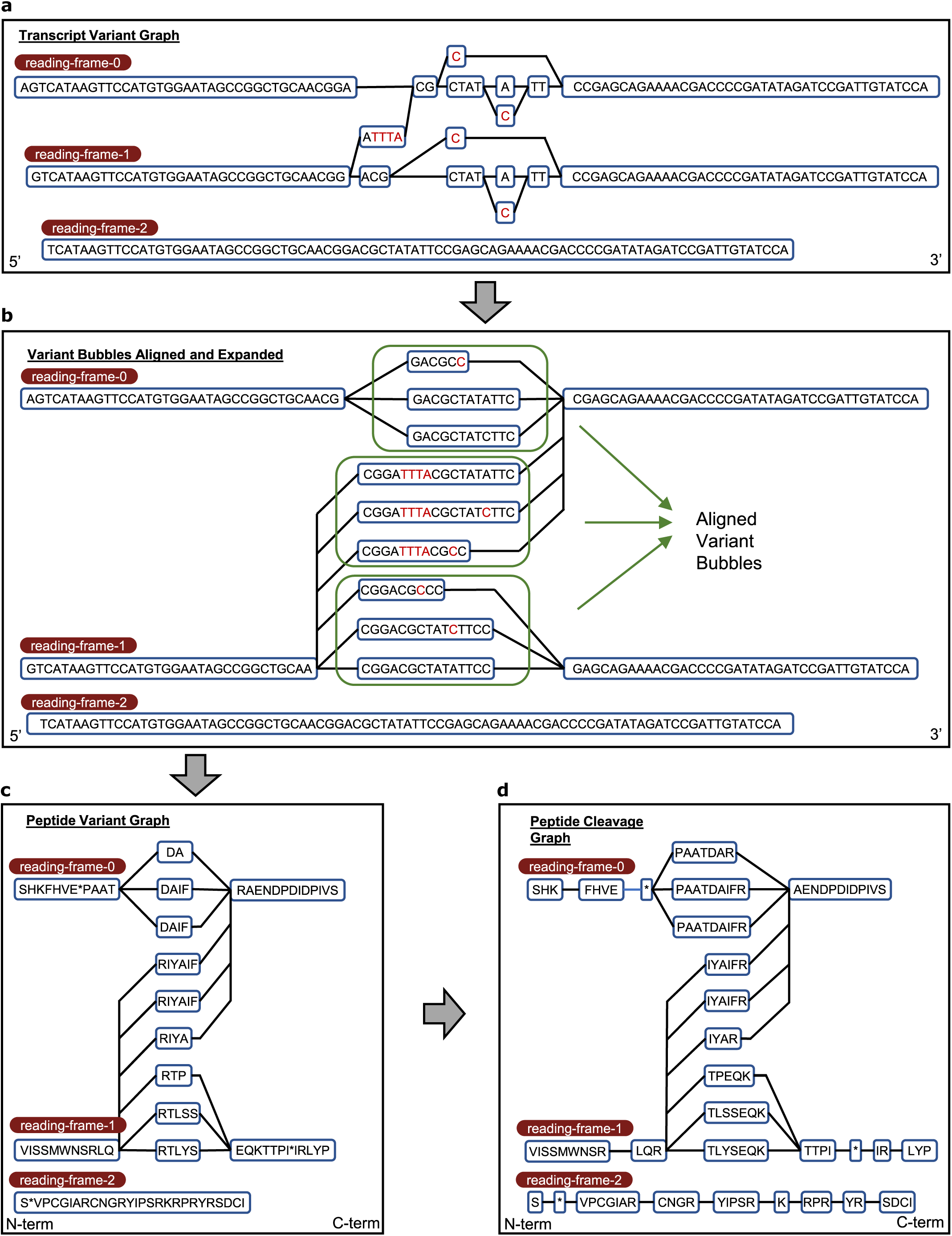
Core graph algorithm of moPepGen. The graph algorithm of moPepGen implements the following key steps: **a**) A transcript variant graph (TVG) is generated from the transcript sequence with all associated variants. All three reading frames are explicitly generated to efficiently handle frameshift variants. **b**) Variant bubbles of the TVG are aligned and expanded to ensure the sequence length of each node is a multiple of three. **c**) Peptide variant graph (PVG) is generated by translating the sequence of each node of the TVG. **d**) Peptide cleavage graph is generated from the PVG in such a way that each node is an enzymatically cleaved peptide.

**Extended Data Figure 2:**
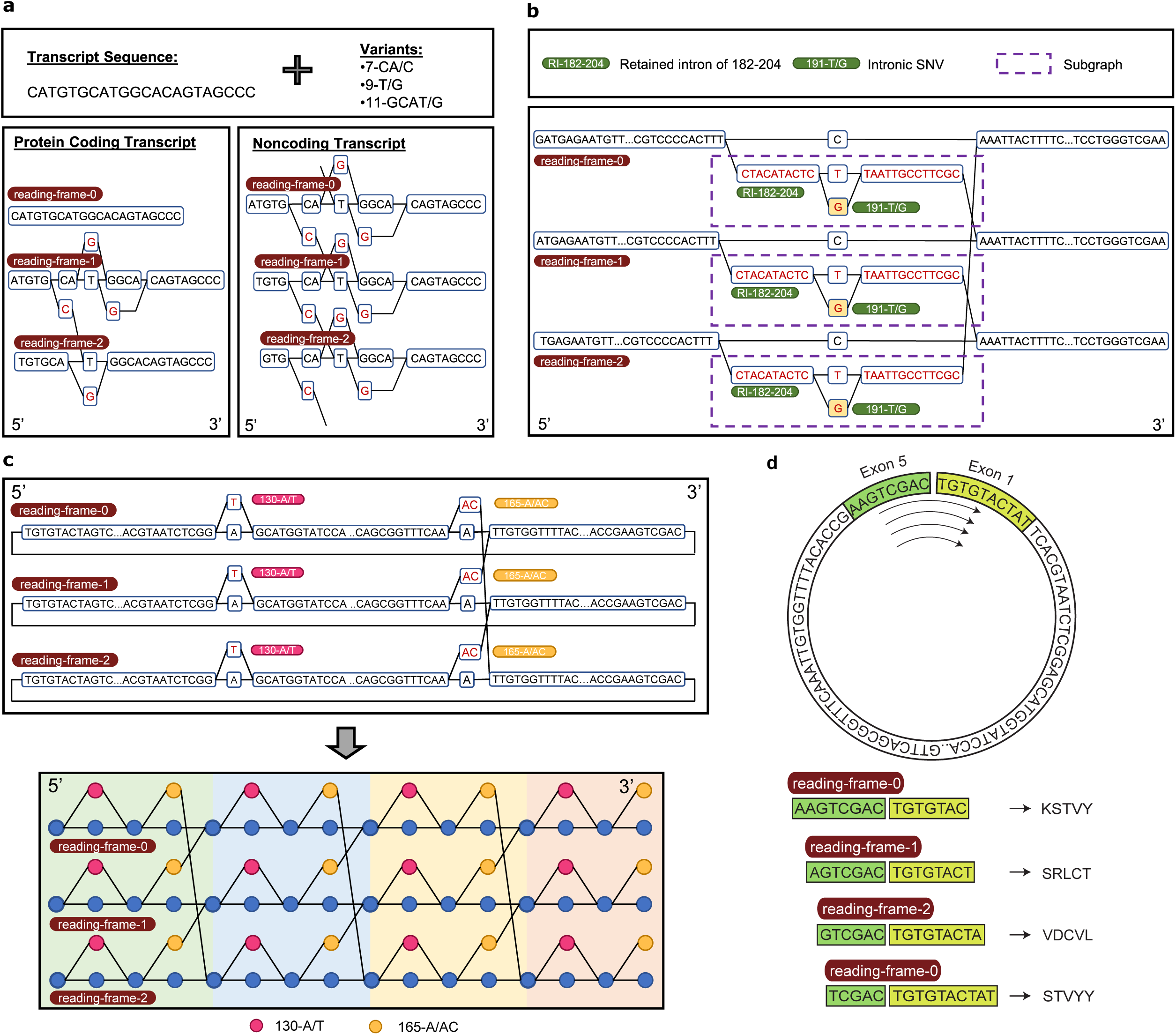
Differential handling of noncoding transcripts, subgraphs and circular RNAs. **a**) For coding transcripts, variants are only incorporated into the effective reading frames. For transcripts that are canonically annotated as noncoding, variants are added to all three reading frames to perform comprehensive three-frame translation. **b**) Subgraphs are created for variant types that involve the insertion of large segments of the genome, which can carry additional variants. **c**) The graph of a circular RNA is extended four times to capture all possible peptides that span the back-splicing junction site in all three reading frames. In the bottom panel, the nodes in magenta harbour the variant 130-A/T and the nodes in yellow harbour 165-A/AC. **d**) Illustration of a circRNA molecule with a novel open reading frame. Each translation across the back-splicing site may shift the reading frame. If no stop codon is encountered, the original reading frame is restored after the fourth crossing.

**Extended Data Figure 3:**
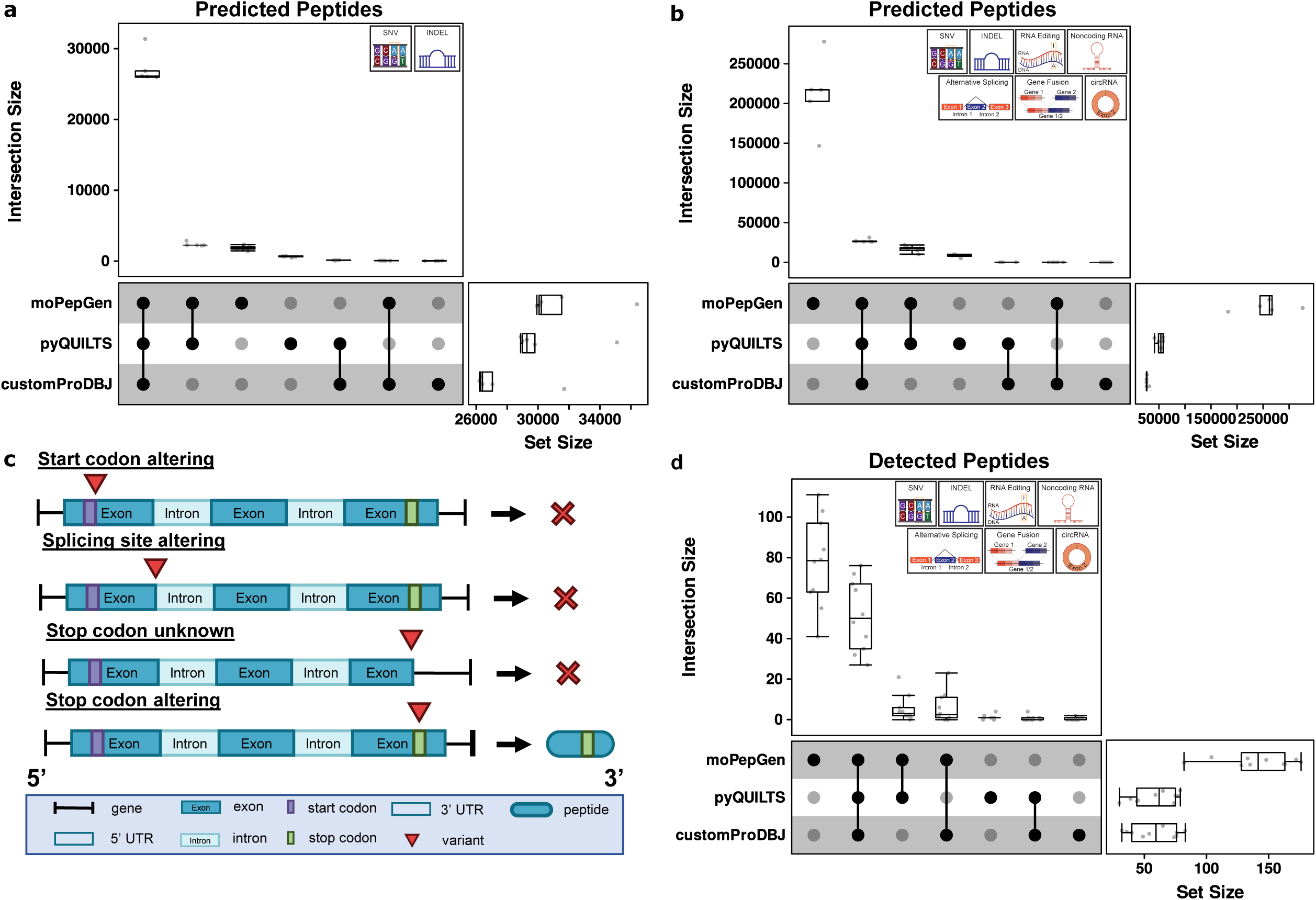
moPepGen demonstrates comprehensive results and deliberate biological assumptions. **a**) and **b**) Non-canonical peptide generation results from benchmarking of moPepGen, pyQUILTS and customProDBJ using only point mutations (SNVs) and small insertions and deletions (indels; **a**), and with inputs from point mutations, indels, RNA editing, transcript fusion, alternative splicing and circular RNAs (circRNAs). **b**). Top boxplot shows the number of peptides in each set intersection and right barplot shows the total number of non-canonical peptides generated by each algorithm in five primary prostate tumour samples (n = 5). **c**) Assumptions made by moPepGen for handling edge cases that differ from other algorithms. Start-codon-altering and splice-site-altering variants are omitted due to the uncertainty of the resulting translation and splicing outcomes. Transcripts with unknown stop codons do not have trailing peptide outputs because of the uncertainty of the trailing enzymatic cleavage site. Stop-codon-altering variants do not result in translation beyond the transcript end, adhering to central dogma. UTR: untranslated region. **d**) Non-canonical database search results from benchmarking of moPepGen, pyQUILTS and customProDBJ using point mutations, indels, RNA editing, transcript fusion, alternative splicing and circRNAs (n = 5). All boxplots show the first quartile, median, to the third quartile, with whiskers extending to furthest points within 1.5× the interquartile range.

**Extended Data Figure 4:**
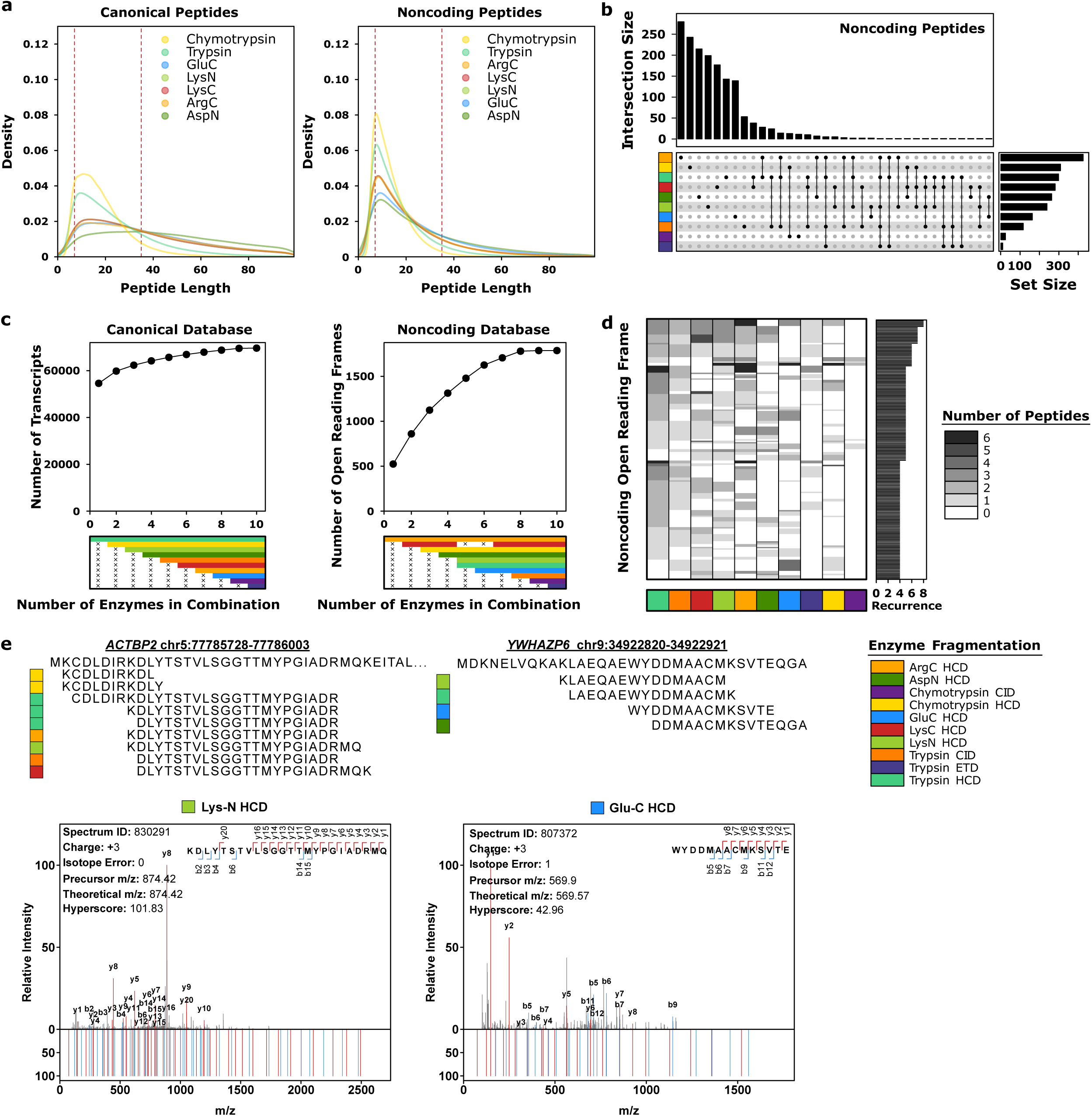
Detection of novel open reading frame peptides across proteases. **a)** Peptide length distributions after *in silico* digestion with seven enzymes, as indicated by color, of the canonical human proteome and three-frame translated noncoding transcript open reading frames (ORFs). The dotted lines indicate the 7-35 amino acids peptide length range commonly used for database search. **b**) Noncoding peptide detection across ten enzyme-fragmentation methods in one deeply fractionated human tonsil sample. The top barplot shows the number of peptides in each set intersection and the right barplot shows the total number of non-canonical peptides from noncoding ORFs detected in each enzyme-fragmentation method, as indicated by covariate color. **c**) Optimal combinations of one to ten enzyme-fragmentation methods for maximizing the number of transcripts detected from the canonical proteome, or the number of ORFs detected from noncoding transcripts. The bottom covariate indicates the optimal combinations of enzyme-fragmentation methods from combinations of one to ten, with color indicating enzyme-fragmentation method. **d**) Noncoding transcript ORFs with peptides detected across four or more enzyme-fragmentation methods, with recurrence count shown in the right barplot. The color of the heatmap indicates the number of peptides detected per ORF per enzyme-fragmentation method. **e**) Example ORFs with coverage by multiple proteases are shown, with peptides tiled according to detection in each enzyme-fragmentation method, as indicated by covariate color. Representative fragment ion mass spectra of peptide-spectrum matches are shown, with theoretical spectra at the bottom and fragment ion matches colored (blue: b-ions, red: y-ions in). HCD: higher-energy collisional dissociation; CID: collision-induced dissociation; ETD: electron-transfer dissociation; *m/z*: mass-to-charge ratio.

**Extended Data Figure 5:**
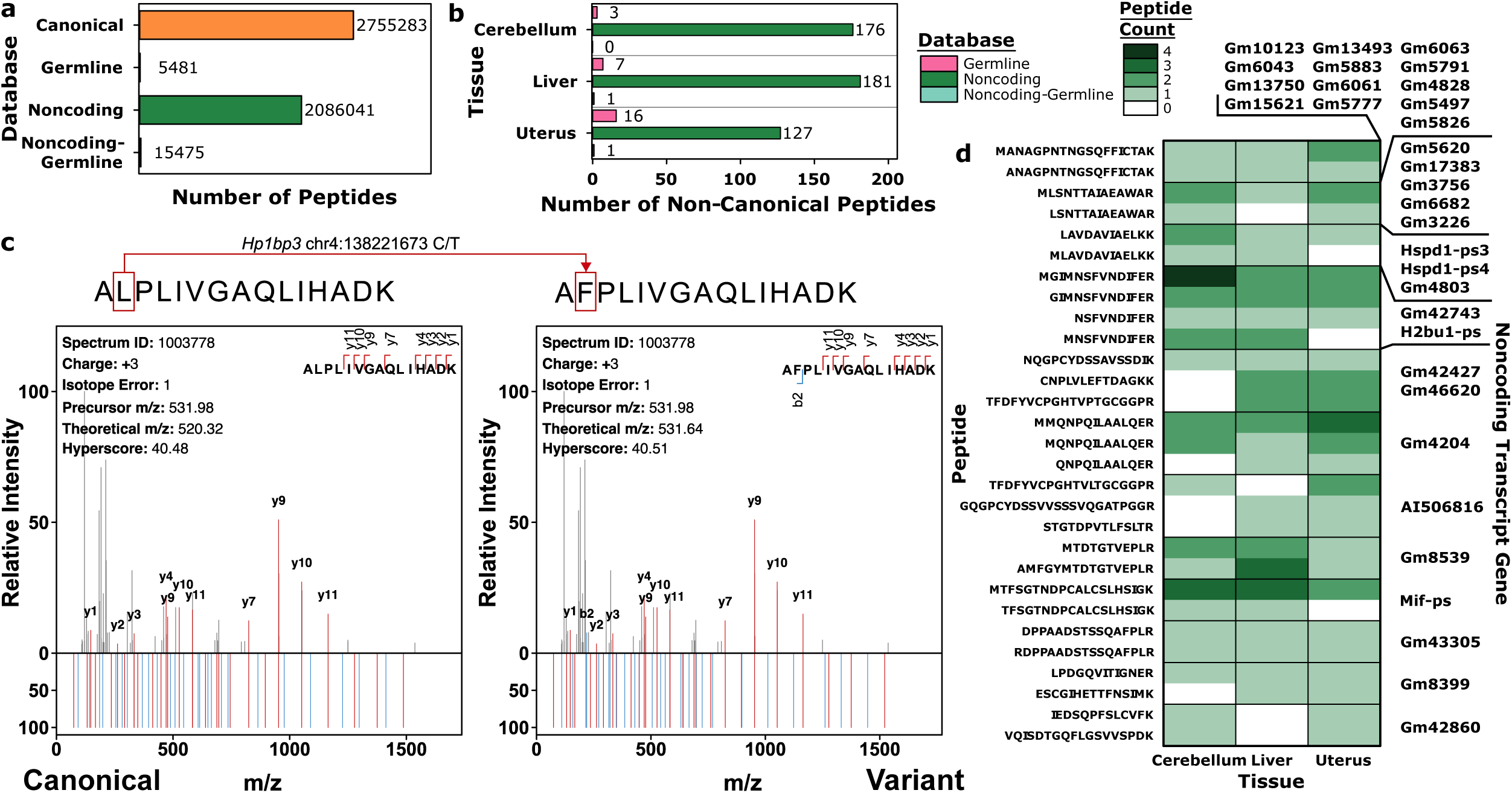
Germline non-canonical peptide detection in mouse strain C57BL/6N. **a)** Comparison of canonical and custom database sizes for the C57NL/6N mouse. Germline database includes single nucleotide polymorphisms (SNPs) and small insertions and deletions. **b)** Number of non-canonical peptides detected from each database in each tissue (one sample per tissue), with database indicated by color. **c**) Comparison of a variant peptide-spectrum match (PSM) spectra (top, both) with the theoretical spectra of the canonical peptide counterpart (left, bottom) as well as the theoretical spectra of the variant peptide harbouring a SNP (right, bottom). Fragment ion matches are colored, with b-ions in blue and y-ions in red. *m/z*: mass-to-charge ratio. **d**) Noncoding transcripts with open reading frames yielding two or more non-canonical peptides recurrently detected across tissues, with color indicating the number of peptides detected in each tissue.

**Extended Data Figure 6:**
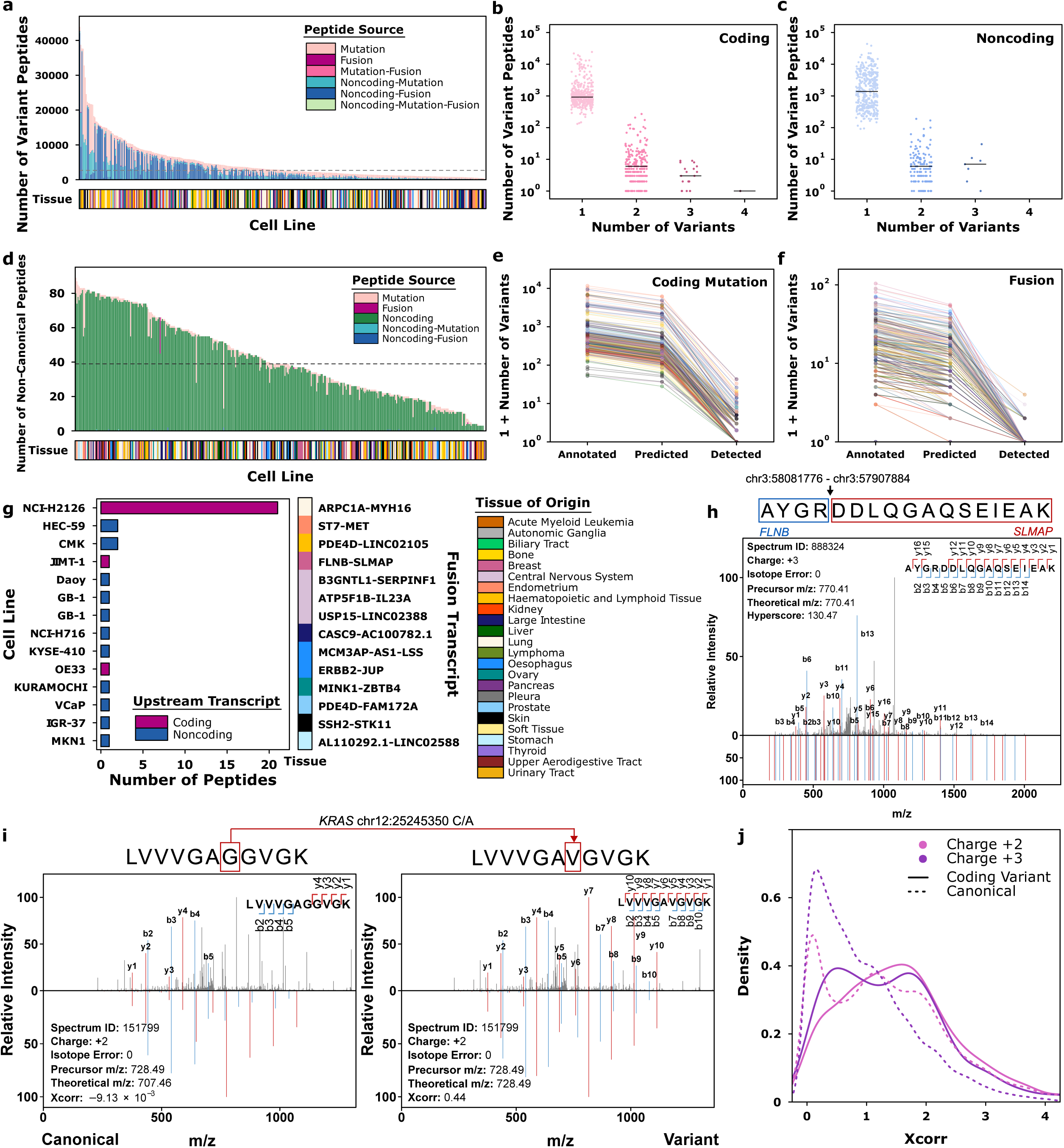
Proteogenomic investigation of the Cancer Cell Line Encyclopedia. **a)** Number of non-canonical peptides generated per cell line, with color indicating peptide source. Bottom covariate indicates tissue of origin. **b**) and **c**) Number of variant peptides per cell line (n = 376) grouped by variant count in coding (**b**) and noncoding (**c**) transcripts. Lines indicate group median. **d**) Number of non-canonical peptides detected per cell line, colored by peptide source. Bottom covariate indicates tissue of origin. **e**) Per cell line, number of intragenic coding mutations (by VEP), mutations predicted to produce detectable non-canonical peptides and mutations detected through proteomics. **f**) Per cell line, number of transcript fusions, those predicted to produce detectable non-canonical peptides and fusions with detected peptide products. Color indicates tissue of origin. **g**) Fusion transcripts (upstream-downstream gene symbol) with detected peptides, with number of peptides shown across cell lines. Bar color indicates whether the upstream fusion transcript was coding or noncoding. Right covariate indicates tissue of origin. **h**) Fragment ion mass spectrum from peptide-spectrum match (PSM) of the non-canonical peptide at the junction of the *FLNB*-*SLMAP* fusion transcript. The peptide theoretical spectrum is shown at the bottom and fragment ion matches are colored (blue: b-ions, red: y-ions). **i**) Comparison of mass spectrum (top, both) from PSM of a non-canonical peptide with a single nucleotide variant against Prosit-predicted MS2 mass spectra based on the canonical counterpart peptide sequence (left, bottom) and the detected variant peptide sequence (right, bottom). Fragment ion matches are colored, with b-ions in blue and y-ions in red. **j**) Cross-correlation (Xcorr) distribution of coding variant peptides PSMs against Prosit-predicted fragment mass spectra (solid lines, color indicate charge), in comparison with Xcorr of control canonical PSMs against Prosit-predicted mass spectra (dotted lines). *m/z*: mass-to-charge ratio.

**Extended Data Figure 7:**
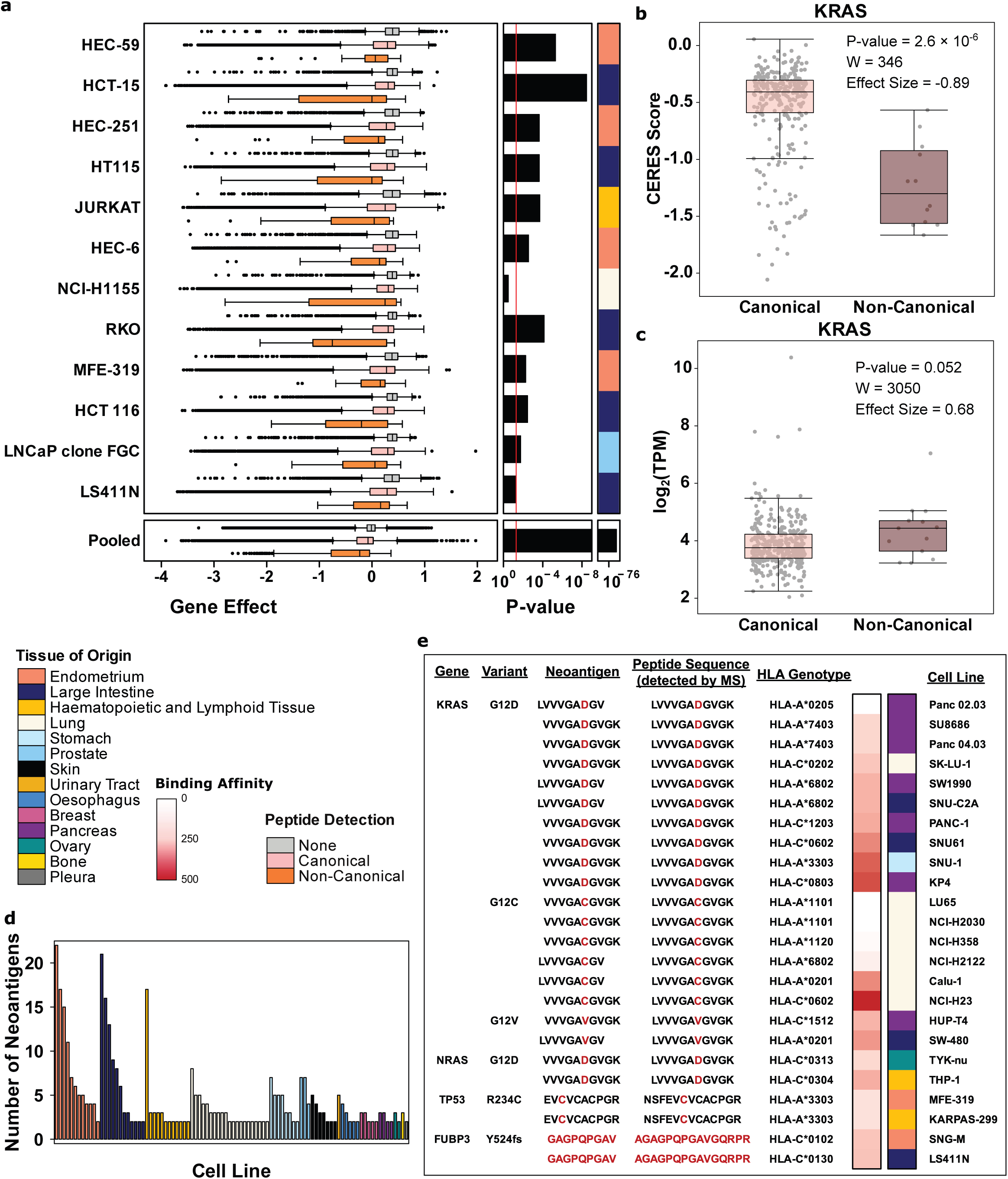
Functional investigation of non-canonical peptide detection in Cancer Cell Line Encyclopedia. **a)** Gene dependency CERES scores for genes with detected non-canonical peptides (orange), detected canonical peptides only (pink) and no detected peptides (gray). A lower CERES score indicates higher gene dependency. Cell lines were selected based on the detection of non-canonical peptides in more than 10 genes. P-values were calculated using a two-sided Mann-Whitney U-test. The red vertical line indicates α = 0.05. The bottom panel represents data pooled across all genes and cell lines. The number of genes per group per cell line and Mann-Whitney U-test results are provided in **Supplementary Table 9**. **b**) Gene dependency CERES score and **c**) mRNA abundance of *KRAS* in cell lines with only canonical peptides detected compared to those with detected non-canonical peptides (n = 290 and 12, respectively). P-values were calculated using a two-sided Mann-Whitney U-test. TPM: Transcript per million. **d**) Number of putative neoantigens predicted based on detected non-canonical peptides in cell lines with more than two neoantigens. The color indicates cell line tissue of origin. **e**) Recurrent neoantigens observed across multiple cell lines, along with their associated gene, variant, *HLA* genotype and the full peptide sequence as detected by trypsin-digested whole cell lysate mass spectrometry. The color in the left heatmap represents neoantigen binding affinity. Right covariate indicates tissue of origin. All boxplots show the first quartile, median, to the third quartile, with whiskers extending to furthest points within 1.5× the interquartile range.

**Extended Data Figure 8:**
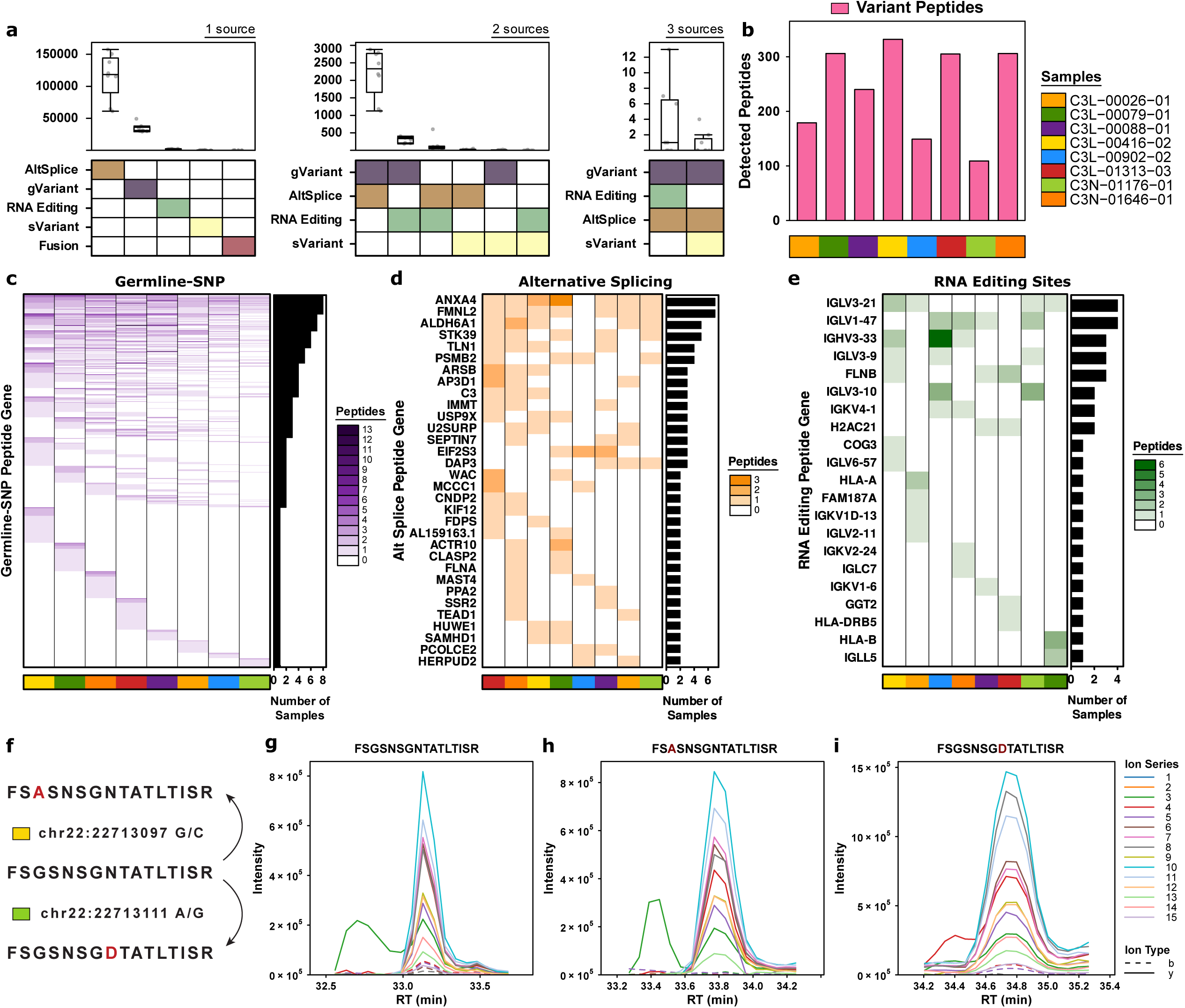
Detection of non-canonical peptides from DIA proteomics. **a)** Number of variant peptides from different variant combinations generated using genomic and transcriptomic data from eight clear cell renal cell carcinoma (ccRCC) tumours (n = 8), grouped by the number of variant sources in combination. gVariant: germline single nucleotide polymorphism and insertion/deletions (indels); sVariant: somatic single nucleotide variant and indels; AltSplice: alternative splicing. **b**) Number of detected variant peptides in the data-independent acquisition (DIA) proteome of eight ccRCC tumours. **c**-**e**) Detection of non-canonical peptides harbouring germline single nucleotide polymorphisms (**c**), alternative splicing (**d**) and RNA editing sites (**e**) across genes. Heatmap colors indicate the number of peptides detected per gene per sample. The barplot indicates recurrence across samples. **f**) Illustration of non-canonical peptides derived from the canonical sequence FSGSNSGNTATLTISR in gene *IGLV3-21* caused by RNA editing events. **g-i**) Extracted ion chromatograms of the canonical peptide (**g**) and non-canonical peptides derived from *IGLV3-21* caused by RNA editing events: chr22:22713097 G-to-C (**h**) and chr22:22713111 A-to-G (**i**). All boxplots show the first quartile, median, to the third quartile, with whiskers extending to furthest points within 1.5× the interquartile range.

**Extended Data Figure 9:**
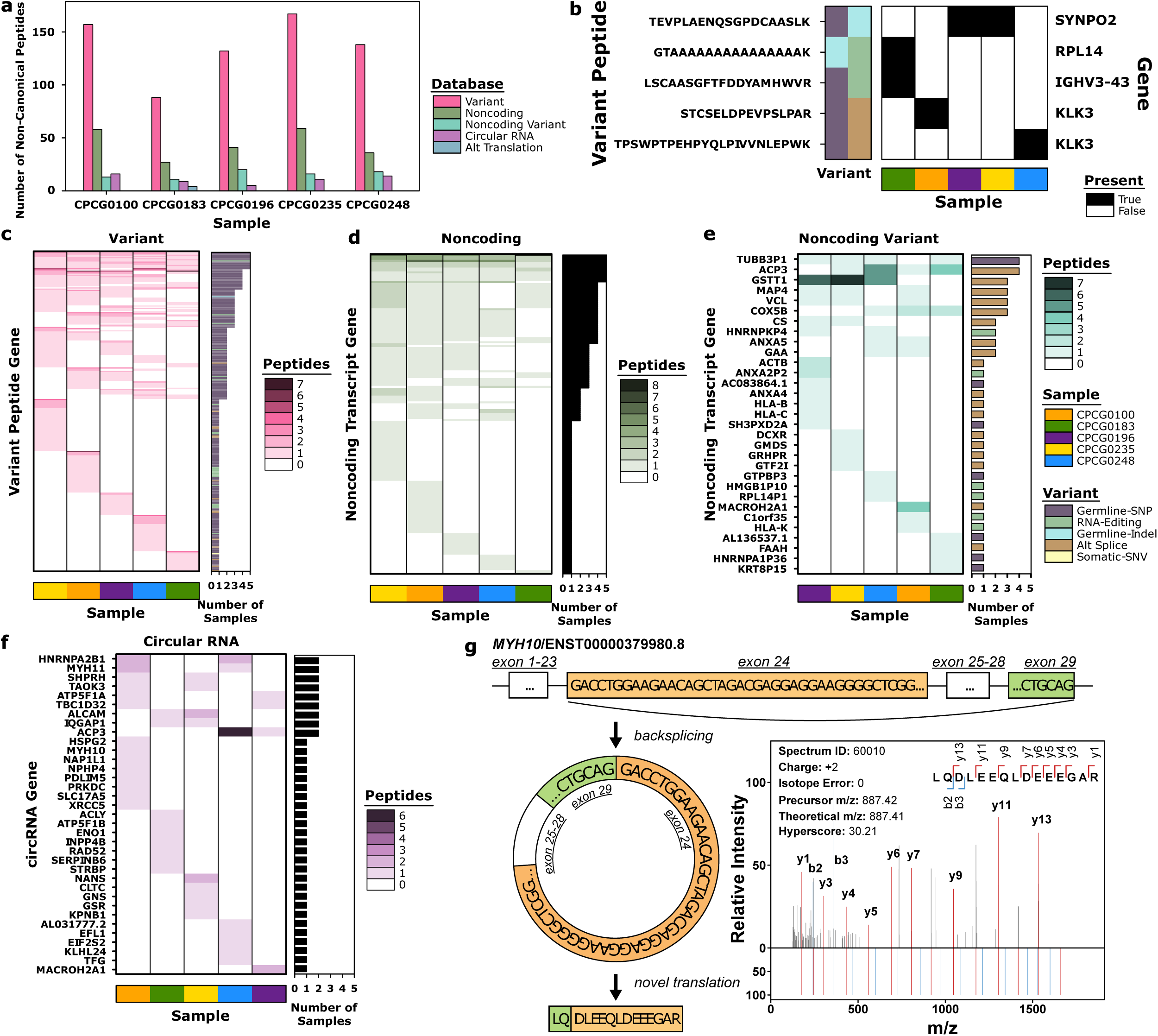
Detection of non-canonical peptides from genomic variants, alternative splicing and circular RNAs. **a)** Number of detected non-canonical peptides in five primary prostate tumour samples per database tier (colored by database). **b**) Peptides as the result of a combination of two variants, with variant type indicated in left covariate and gene on the right. The heatmap shows presence of peptide across samples. **c**-**f**) Non-canonical peptide detection results across genes, with color of heatmap representing the number of peptides detected per gene per sample. The barplot indicates recurrence across samples, and when colored indicates variant type associated with the gene entry. The Variant database includes non-canonical peptides from coding transcripts with single nucleotide polymorphisms (SMPs), single nucleotide variants (SNVs), small insertion and deletion (indels), RNA editing, alternative splicing (Alt Splice) or transcript fusion (**c**). Noncoding database includes all peptides from noncoding transcript three-frame translation open reading frames (**d**) and noncoding peptides with any variants are included in the Noncoding Variant database (**e**). The Circular RNA database includes all peptides representing circular RNA open reading frames (ORFs) with or without other variants (**f**). The bottom covariate indicates prostate cancer sample. **g**) Mass spectrum from peptide-spectrum match of a non-canonical peptide spanning the back-splicing junction between exon 29 and exon 24 of *MYH10*, reflective of circular RNA translation. The peptide theoretical spectrum is shown at the bottom and fragment ion matches are colored (blue: b-ions, red: y-ions in). *m/z*: mass-to-charge ratio.

## Extended Data Figure Legend

**Extended Data Table 1:**
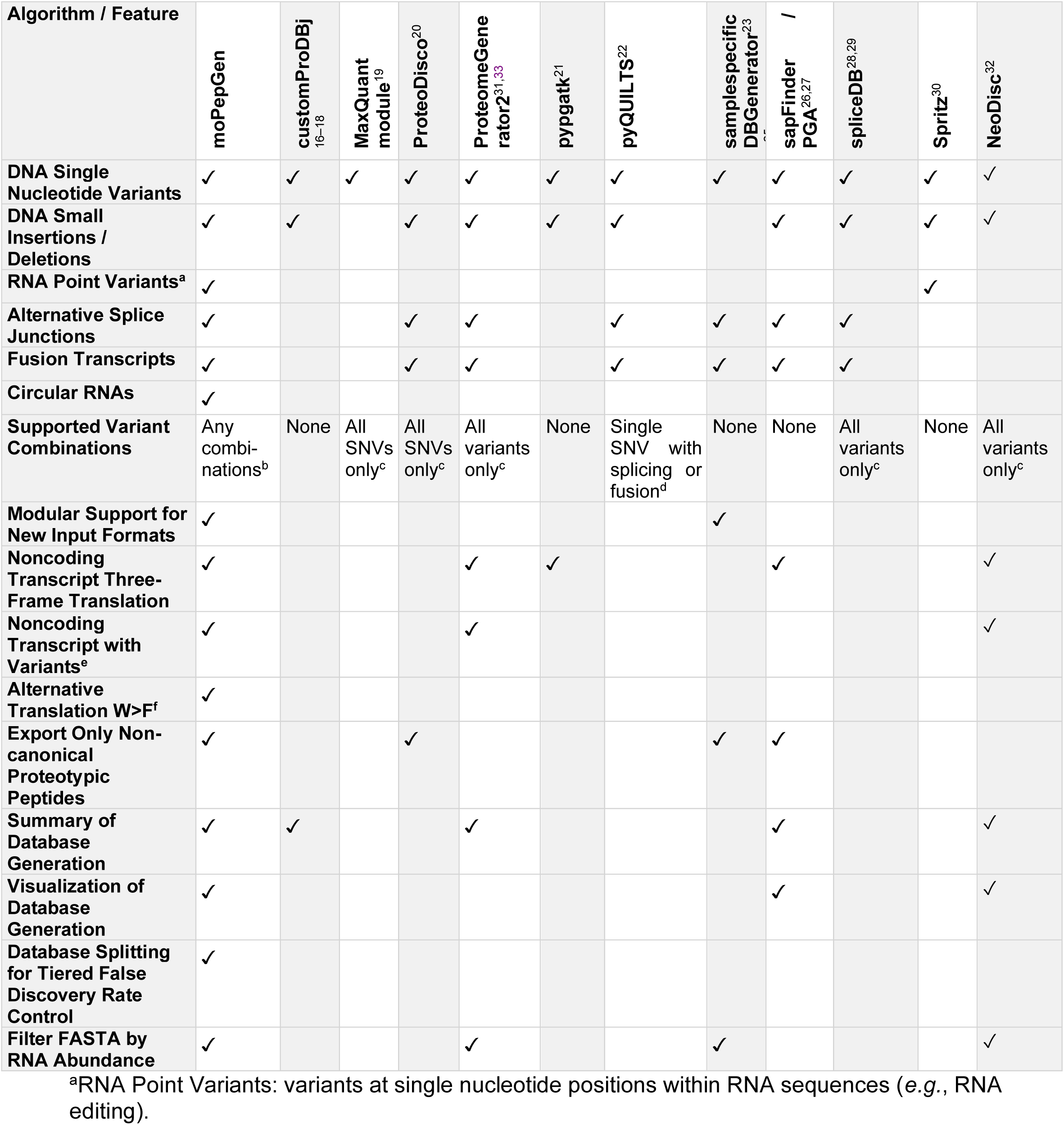
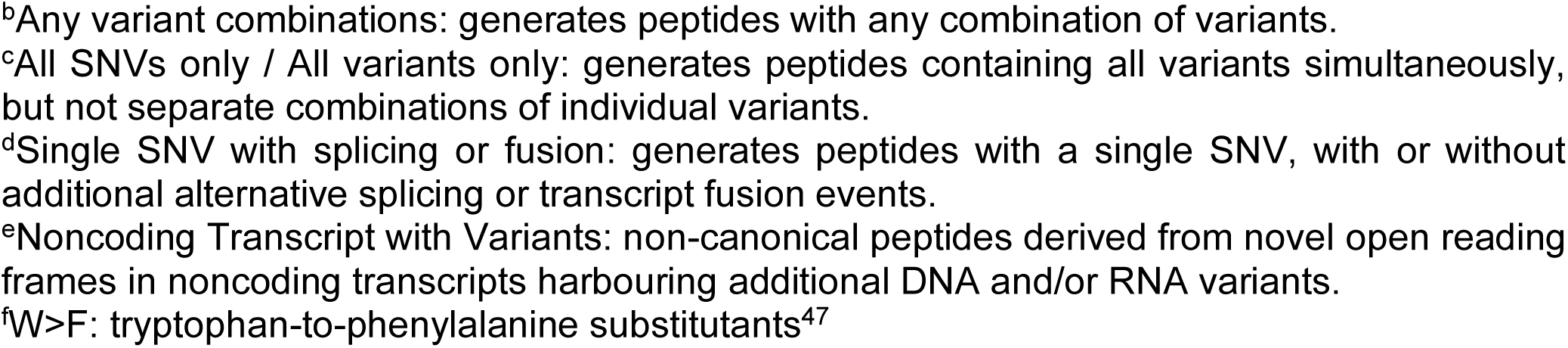
Feature comparison of custom database generation algorithms.

| Algorithm / Feature | moPepGen | customProDBj<sup>16–18</sup> | MaxQuant module<sup>19</sup> | ProteoDisco<sup>20</sup> | ProteomeGenerator<sup>21,33</sup> | pypgatk<sup>21</sup> | pyQUILTS<sup>22</sup> | samplespecific DBGenerator<sup>23</sup> | sapFinder / PGA<sup>26,27</sup> | spliceDB<sup>28,29</sup> | Spritz<sup>30</sup> | NeoDisc<sup>32</sup> |
| --- | --- | --- | --- | --- | --- | --- | --- | --- | --- | --- | --- | --- |
| DNA Single Nucleotide Variants | ✓ | ✓ | ✓ | ✓ | ✓ | ✓ | ✓ | ✓ | ✓ | ✓ | ✓ | ✓ |
| DNA Small Insertions / Deletions | ✓ | ✓ |  | ✓ | ✓ | ✓ | ✓ |  | ✓ | ✓ | ✓ | ✓ |
| RNA Point Variants<sup>a</sup> | ✓ |  |  |  |  |  |  |  |  |  | ✓ |  |
| Alternative Splice Junctions | ✓ |  |  | ✓ | ✓ |  | ✓ | ✓ | ✓ | ✓ |  |  |
| Fusion Transcripts | ✓ |  |  | ✓ | ✓ |  | ✓ | ✓ | ✓ | ✓ |  |  |
| Circular RNAs | ✓ |  |  |  |  |  |  |  |  |  |  |  |
| Supported Variant Combinations | Any combinations<sup>b</sup> | None | All SNVs only<sup>c</sup> | All SNVs only<sup>c</sup> | All variants only<sup>c</sup> | None | Single SNV with splicing or fusion<sup>d</sup> | None | None | All variants only<sup>c</sup> | None | All variants only<sup>c</sup> |
| Modular Support for New Input Formats | ✓ |  |  |  |  |  |  | ✓ |  |  |  |  |
| Noncoding Transcript Three-Frame Translation | ✓ |  |  |  | ✓ | ✓ |  |  | ✓ |  |  | ✓ |
| Noncoding Transcript with Variants<sup>e</sup> | ✓ |  |  |  | ✓ |  |  |  |  |  |  | ✓ |
| Alternative Translation W>F<sup>f</sup> | ✓ |  |  |  |  |  |  |  |  |  |  |  |
| Export Only Non-canonical Proteotypic Peptides | ✓ |  |  | ✓ |  |  |  | ✓ | ✓ |  |  |  |
| Summary of Database Generation | ✓ | ✓ |  |  | ✓ |  |  |  | ✓ |  |  | ✓ |
| Visualization of Database Generation | ✓ |  |  |  |  |  |  |  | ✓ |  |  | ✓ |
| Database Splitting for Tiered False Discovery Rate Control | ✓ |  |  |  |  |  |  |  |  |  |  |  |
| Filter FASTA by RNA Abundance | ✓ |  |  |  | ✓ |  |  | ✓ |  |  |  | ✓ |

<sup>a</sup>RNA Point Variants: variants at single nucleotide positions within RNA sequences (e.g., RNA editing).
<sup>b</sup>Any variant combinations: generates peptides with any combination of variants.
<sup>c</sup>All SNVs only / All variants only: generates peptides containing all variants simultaneously, but not separate combinations of individual variants.
<sup>d</sup>Single SNV with splicing or fusion: generates peptides with a single SNV, with or without additional alternative splicing or transcript fusion events.
<sup>e</sup>Noncoding Transcript with Variants: non-canonical peptides derived from novel open reading frames in noncoding transcripts harbouring additional DNA and/or RNA variants.
<sup>f</sup>W>F: tryptophan-to-phenylalanine substitutants<sup>47</sup>

