## Supplementary Information for "Identification of non-canonical peptides with moPepGen"

#### Supplementary Notes

##### Supplementary Note 1: Terminology

Canonical Database

A *canonical database* is a reference protein database containing amino acid sequences derived exclusively from well-annotated protein-coding transcripts without variants or modifications. It represents the standard, widely accepted protein sequences in a given organism.

Non-canonical Peptides

Non-canonical peptides are absent from the canonical database. They can arise from genomic variants (*e.g.*, germline single nucleotide polymorphisms (SNPs), somatic single nucleotide variants (SNVs), small insertion/deletions (indels)), transcriptomic events (*e.g.*, alternative splicing, transcript fusion, RNA editing, circRNAs), and sequence modifications during translation (*e.g.*, selenocysteine terminations, tryptophan-to-phenylalanine (W>F) substitutants) in protein-coding transcripts. Peptides translated from novel open reading frames (ORFs) in transcripts annotated as noncoding (*e.g.*, lncRNAs, pseudogenes) are also classified as non-canonical peptides when derived from valid biological processes, with possible further additions of genomic or transcriptomic variants.

Variant Peptides

Variant peptides are a subset of non-canonical peptides harbouring genomic and/or transcriptomic variants translated from protein-coding transcripts.

Non-canonical Database

A *non-canonical database* contains peptide or protein amino acid sequences derived from genomic or transcriptomic variants, novel alternative splicing events, or previously unannotated and misannotated non-coding regions. By ensuring no overlap with the canonical database, this database enables the detection of unique, rare, or condition-specific peptides that cannot be detected using canonical databases alone.

Proteoform

A proteoform is a specific molecular form of a protein with a unique amino acid sequence resulting from genetic variations (*e.g.*, germline SNPs, somatic SNVs, indels), transcriptional processes (*e.g.*, alternative splicing, transcript fusion, RNA editing), translation-level sequence modifications (*e.g.*, selenocysteine terminations, W>F substitutants) or post-translational/co-translational modifications generated by enzymatic (*e.g.*, phosphorylation, acetylation, N/O-glycosylation) or non-enzymatic (*e.g.*, glycation, oxidation, etc.) attachment of specific chemical moieties to the side chains of specific amino acids.

**Supplementary Note 2**: Finding the connection node that serves as the commonly connected downstream node of a variant bubble.

---

**Algorithm 1**: Find the next connection node after the variant bubble

---

**Input**: connection node $n$, minimal connection node length $j$
**Output**: next connection node

```
Function FindNextConnectionNode(n, j) begin
    Initialize Q as empty queue, n_t as null
    For n_o in n.OutNodes
        Append n_o to Q
    While Q is not empty
        cur ← Q.popLast()
        If cur has no variants and (n_t = null or cur > n_t) and length(cur) ≥ j
            n_t ← cur
        else
            For n_i in cur.OutNodes
                Append n_i to Q
    Return n_t
```

---

**Supplementary Note 3:** Assigning ORF to nodes using the *stage-and-call* approach.

**Algorithm 2:** Call Variant Peptides using the stage-and-call approach

```
Input: root node r, ORF start site s, allowed miscleavages k
Output: Set of variant peptides

Function StageAndCall(r, s, k) begin
    Initialize Q as empty queue, V as empty hash table, P as empty set
    For n in r.OutNodes:
        Append [n, False] to Q
    While Q is not empty
        [n_i, l] ← Q.popFirst()
        If l = True
            For p in n_i.GetMiscPeptides(k)
                Add p to P
        Else If s in n_i
            l ← True
            n_i' ← n_i.RemoveSequenceLeftTo(s)
            For p in n_i'.GetMiscPeptides(k)
                Add p to P
        For n_o in n_i.OutNodes
            Append [n_i, l] to V[n_o]
            If length(V[n_o]) = n_o.InNodes:
                l_o ← any l = True for [n, l] in V[n_o]
                Append [n_o, l_o] to Q
    Return P
```

**Supplementary Note 4:** Identifying variant peptides with permitted number of miscleavages.

---

**Algorithm 3:** Get miscleaved peptide sequences

---

**Input:** Peptide cleavage graph node $n$, maximal allowed miscleavages $k$
**Output:** Set of variant peptides

**Function** GetMiscPeptides($n$, $k$) **begin**
  **Initialize** $P$ as empty set, $Q$ as empty queue
  **Append** [$n$] to $Q$
  **While** $Q$ is not empty
    $N \leftarrow Q$.pop()
    **If** any $n_i$.variants is not empty for $n_i$ in $N$:
      Set $p$ to empty string
      **For** $n_i$ **in** $N$
        $p$ += $n_i$.sequence
      **Append** $p$ to $P$
    **If** length($N$) – 1 < $k$
      **For** $n_o$ in $N$[-1].OutNodes:
        $N' \leftarrow N$ + [$n_o$]
        **Append** $N'$ to $Q$
  **Return** $P$

---

### Supplementary Figures

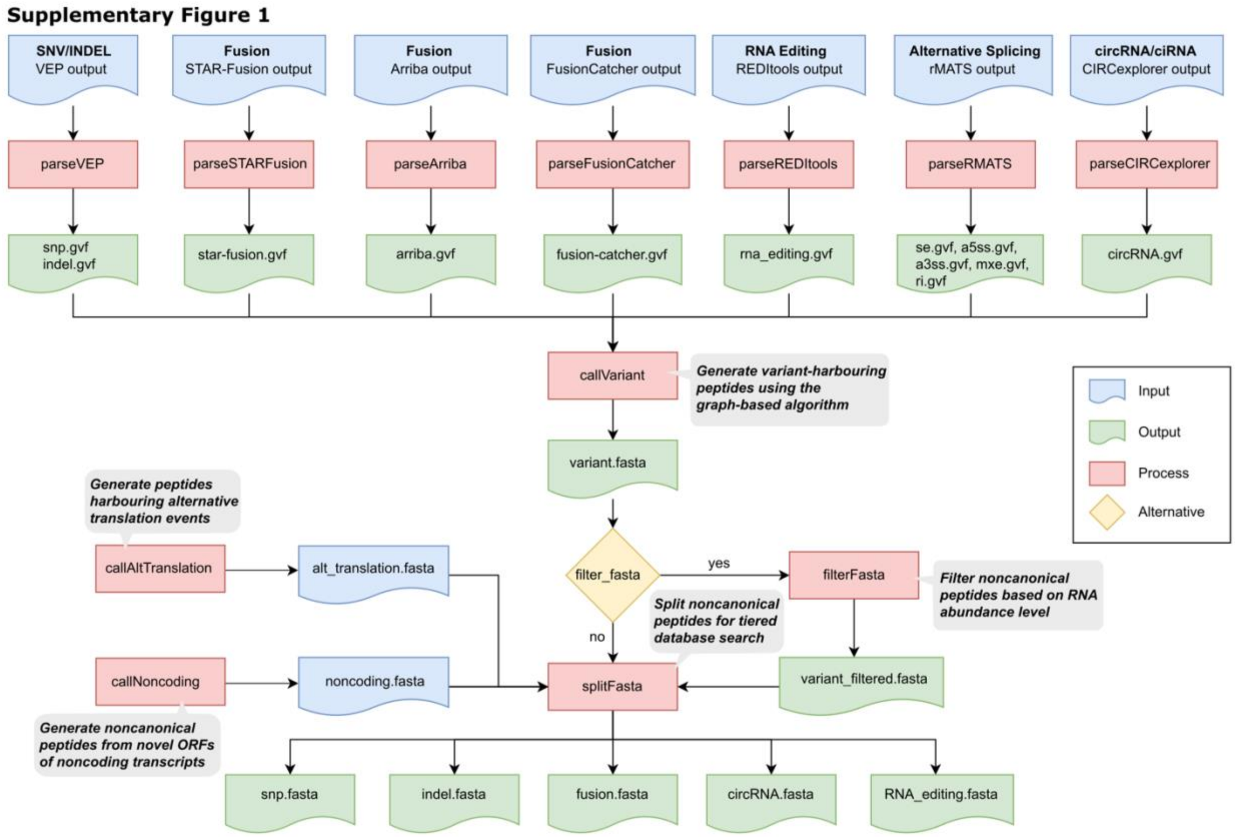


#### *Supplementary Figure 1: Custom Database Generation Pipeline*

Inputs from variant callers are first processed through dedicated parsing modules as part of moPepGen, generating inputs to moPepGen in the form of GVF files. Non-canonical databases are called by the callVariant module, representing the central functionality of moPepGen. Non-canonical peptides from novel open reading frames and alternative translation are called through separate modules and could merge with variant peptides and be split into individual databases for tiered custom database searching. Variant peptides could additionally be filtered by transcript expression levels.

Supplementary Figure 2

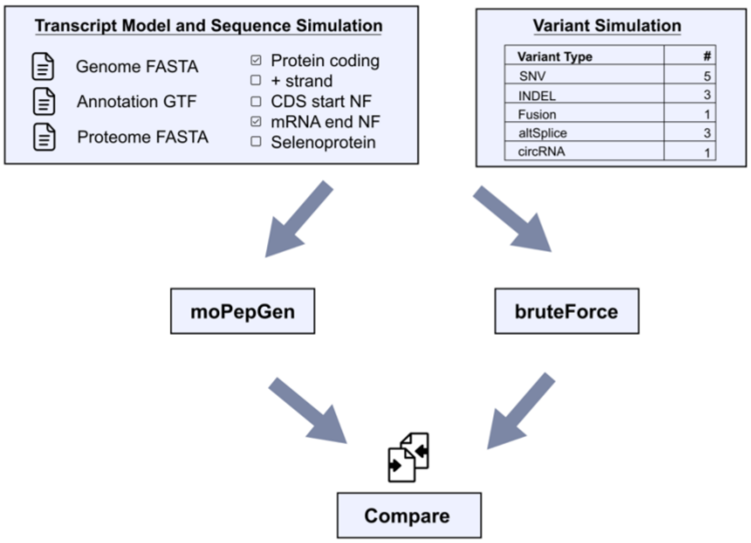


***Supplementary Figure 2: Fuzz testing schema***

In each fuzz test scenario, a simulated transcript annotation model and nucleotide sequence are generated, accompanied by a comprehensive set of variant records encompassing all variant types compatible with moPepGen. Variant peptides are then generated using moPepGen, and the results are validated with those obtained from a brute-force algorithm. This algorithm identifies non-canonical peptides by exhaustively iterating through every potential combination of variants.

**Supplementary Figure 3**

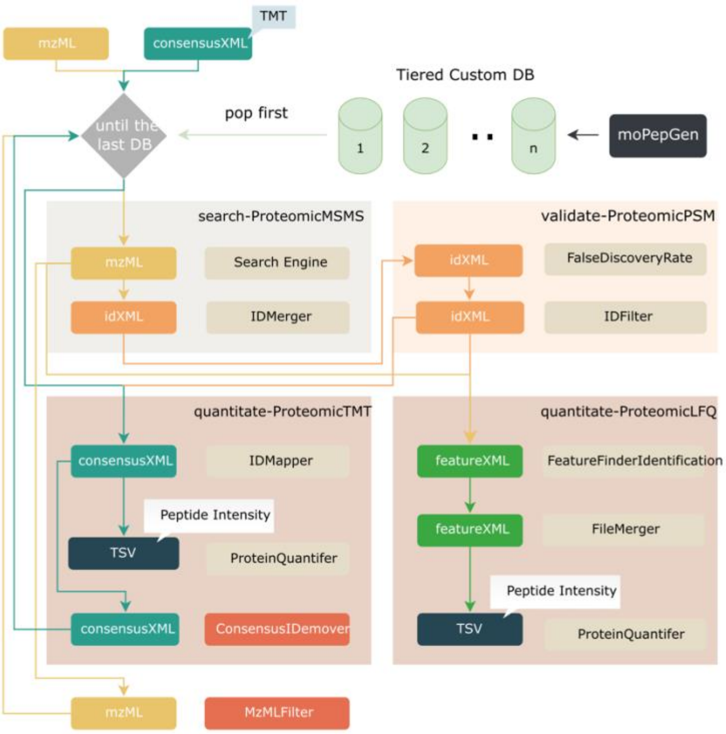


#### ***Supplementary Figure 3: Tiered database search schema***

Individual pipelines and modules involved in database searching against custom databases. Databases are searched in a tiered system where peptide spectrum matches with false-discovery rate (FDR) of 1% at the peptide level are removed from the mzML before searching against the next database, for a conservative approach to non-canonical peptide FDR control.

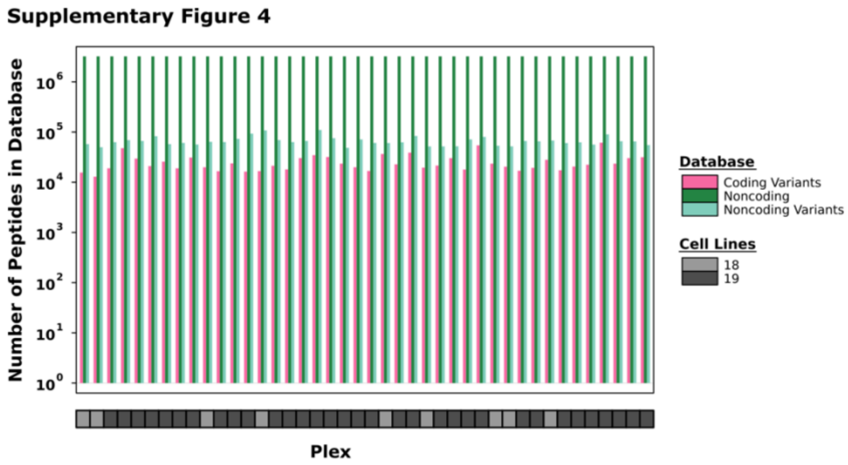


##### *Supplementary Figure 4: Merged database sizes per plex*

Sizes of plex-level custom databases used for non-canonical peptide search in Cancer Cell Line Encyclopedia (CCLE) tandem mass tag (TMT) proteomics data. Each plex database was generated by first merging all variant peptide databases from all cell lines of the plex, including the ten cell lines in the reference channel, followed by splitting into peptides with variants on coding transcripts, peptides from novel open reading frames and peptides from variants on noncoding transcripts, as indicated by color. Bottom covariate indicates the number of unique cell lines in each of the 42 plex.

**Supplementary Figure 5**

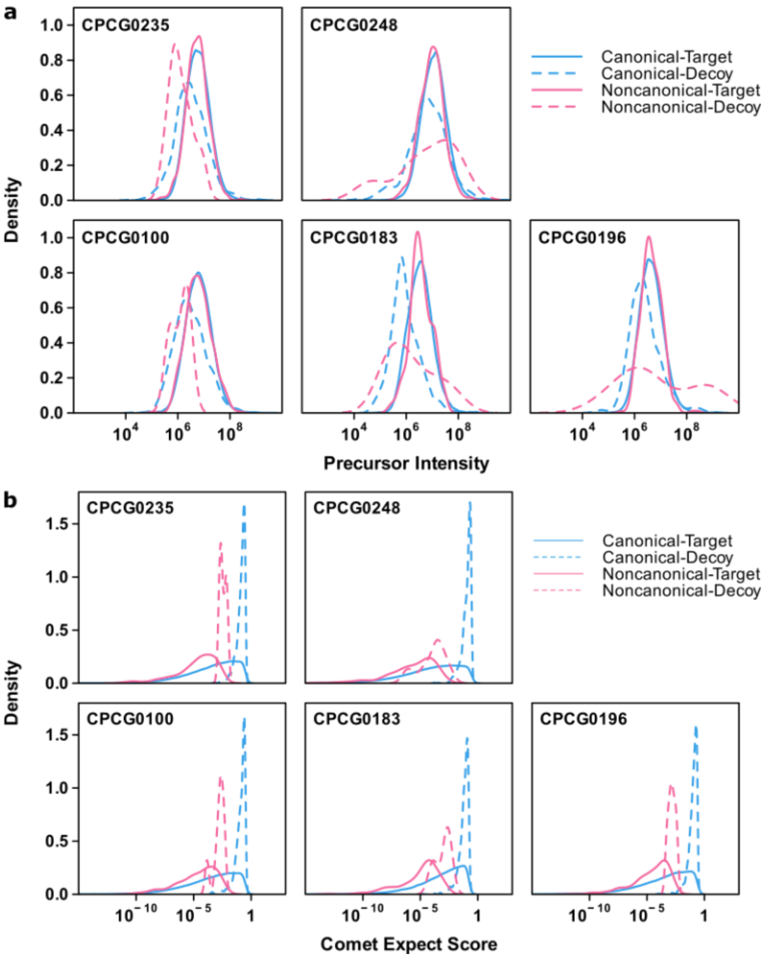


#### *Supplementary Figure 5: Precursor Intensities and Comet Expectation Scores*

Precursor intensities (**a**) and Comet expectation scores (**b**) for canonical and non-canonical PSMs, distinguishing between target and decoy peptides, across five prostate tumour samples.

**Supplementary Figure 6**

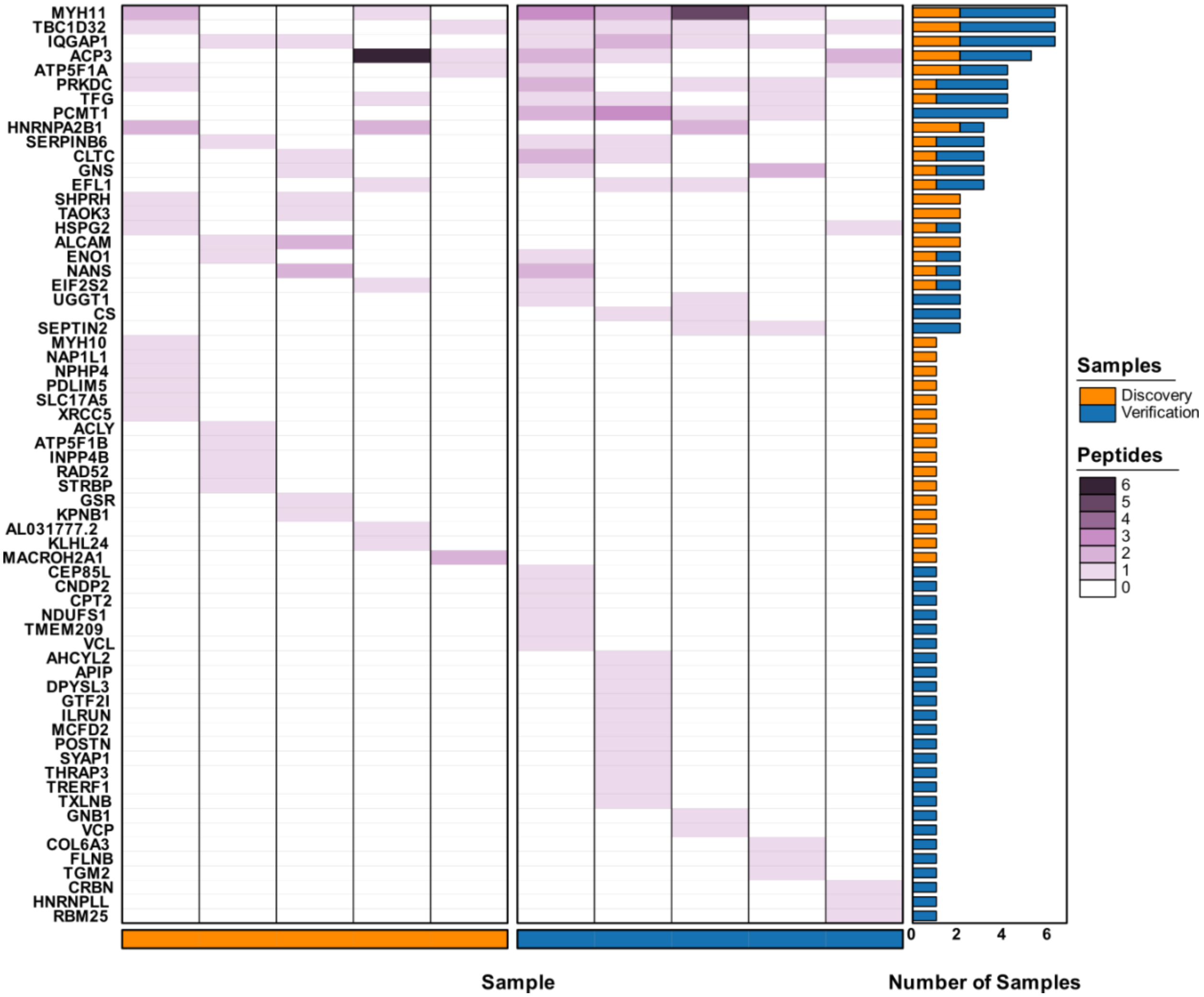


***Supplementary Figure 6: Detection of Non-Canonical Peptides from circular RNAs***

**Supplementary Figure 5**: Database search results for non-canonical peptides derived from circular RNA molecules in discovery (left) and verification (right) samples. Verification samples were selected based on the similarity of circular RNA junction counts with the discovery samples. The color of heatmap represents the number of peptides detected per gene per sample. The barplot indicates recurrence across samples, with discovery (orange) and verification (blue) set separated in colors.

**Supplementary Figure 7**

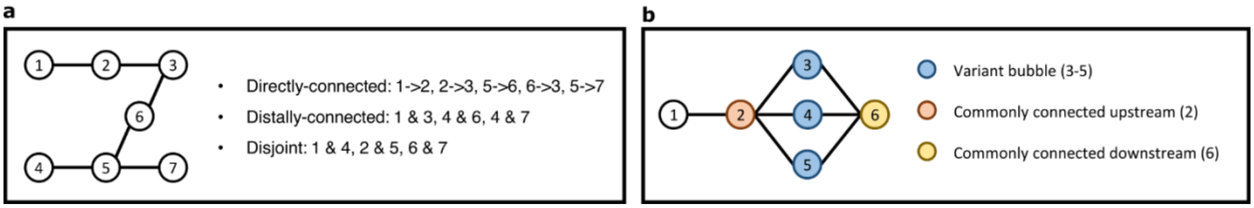


#### *Supplementary Figure 7: Node relationships*

**a**) Examples of nodes that are directly and distally connected, and nodes that are disjoint.

**b**) A variant bubble is defined as a group of connected nodes sharing the same commonly connected upstream (node 2) and downstream (node 6). In this example, node 3, 4, and 5 are members of the variant bubble.

**Supplementary Figure 8**

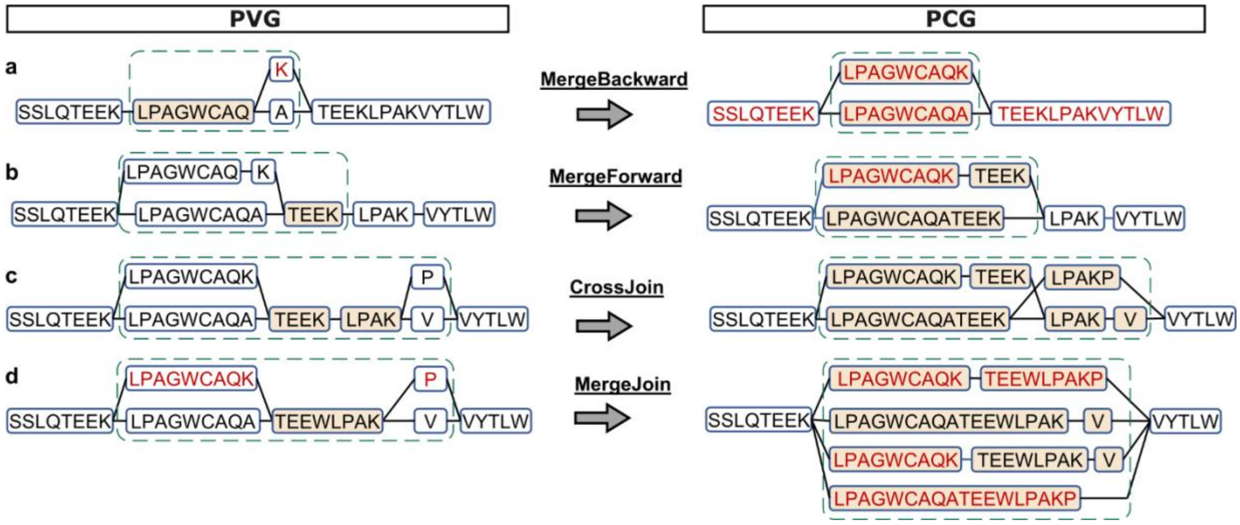


#### *Supplementary Figure 8: Merge and cleave*

Various "merge-and-cleave" approaches are employed to generate a peptide cleavage graph (PCG, right) from a peptide variant graph (PVG, left). **a**) "Merge-backward" is used when a node has a single incoming node and multiple outgoing nodes, and the incoming node represents an enzymatically cleaved peptide. The multiple outgoing nodes are merged with it and the resulting nodes are cleaved at any cleavage sites. **b**) "Merge-forward" is employed when a node has multiple incoming nodes and a single outgoing node. Similar to "merge-backward", incoming nodes are merged with it, followed by enzymatic cleavage. **c**) "Cross-join" is used when the upstream node of two exclusively connected nodes has multiple incoming nodes, and the downstream node also has multiple outgoing nodes. All incoming nodes are merged with the upstream and all outgoing nodes are merged with the downstream node. All possible connection edges are created between the merged upstream and downstream nodes. All merged nodes are also cleaved at any cleavage sites. **d**) "Merge-join" is used when a node has multiple incoming and outgoing nodes. All combinations of incoming and outgoing nodes are merged and cleaved at any cleavage sites.

### Supplementary Tables

#### *Supplementary Table 1: Alternative Protease Non-Canonical Peptides*

Non-canonical peptide detection and annotation results for tonsil sample processed independently with ten protease and fragmentation method combinations. Detected peptides with modifications, charges and intensities are shown for each protease and fragmentation method combination, and annotated with FASTA headers from moPepGen. Sources of each non-canonical peptide are designated for each FASTA header and for the peptide overall.

#### *Supplementary Table 2: Mouse Proteome Non-Canonical Peptides*

Non-canonical peptide detection and annotation results for the liver, uterus and cerebellum proteomes of mouse strain C57BL/6N. Detected peptides with modifications, charges and intensities are shown for each tissue, and annotated with FASTA headers from moPepGen. Sources of each non-canonical peptide are designated for each FASTA header and for the peptide overall.

#### *Supplementary Table 3: Cancer Cell Line Encyclopedia Non-Canonical Peptides*

Non-canonical peptide detection and annotation results for 375 cell line proteomes from the Cancer Cell Line Encyclopedia (CCLE). Detected peptides with modifications, charges and intensities are shown for each cell line and plex, and annotated with FASTA headers from moPepGen. Sources of each non-canonical peptide are designated for each FASTA header and for the peptide overall.

#### *Supplementary Table 4: Neoantigens Predicted from Detected Non-Canonical Peptides.*

Putative neoantigens predicted from detected non-canonical peptides derived from somatic mutations. Neoantigenic peptides with flanking amino acids are shown for each cell line and plex, along with MHC binding affinities based on cell-line-specific *HLA* genotypes.

##### *Supplementary Table 5: DIA Proteome Non-canonical Peptides*

Non-canonical peptide detection results for eight data-independent acquisition (DIA) proteomes. Detected peptides are shown with modifications, charges, intensities, retention times, and statistical scores estimated by DIA-NN for each sample. Sources of each non-canonical peptide are designated for each FASTA header and for the peptide overall.

##### *Supplementary Table 6: Prostate Cancer Non-Canonical Peptides*

Non-canonical peptide detection and annotation results for 5 prostate cancer proteomes. Per database tier, detected peptides with modifications, charges and intensities are shown for each sample, and annotated with FASTA headers from moPepGen. Sources of each non-canonical peptide are designated for each FASTA header and for the peptide overall.

##### *Supplementary Table 7: Circular RNA Novor Validation*

*De novo* sequencing results from Rapid Novor on spectra of peptide spectrum matches (PSMs) against circular RNA peptides for the five discovery and five verification prostate cancer samples. PSM information includes charge, spectrum ID, precursor ion *m/z*, isotope error, retention time, detected circular RNA peptide sequence, theoretical peptide *m/z* and *q*-value. *De novo* sequencing peptide sequence is included with Novor score, *de novo* mass error and whether the *de novo* sequence matched with the PSM.

##### *Supplementary Table 8: Number of cell lines per tissue of origin included in this study*

Number of cancer cell lines analyzed in this study, categorized by tissue of origin. Inclusion was based on availability of both proteomic and genomic data.

##### *Supplementary Table 9: Gene dependency associations with non-canonical peptide detection.*

For each of the 12 cell lines with the most non-canonical peptides detected, genes were categorized based on detection of non-canonical, canonical-only, or no peptides in proteomics data. Median and MAD of CERES gene dependency scores are reported per group. Gene dependency differences between groups were tested using a two-sided Mann–Whitney U test. The pooled row aggregates genes across all 12 cell lines.

#### Supplementary Data

##### *Supplementary Data: Cancer Cell Line Variant Peptide Database*

Custom peptide database FASTA file for each of the 376 cell lines from the cancer cell line encyclopedia with proteomics characterization or used in the reference channel. Variant peptide FASTA files were produced by moPepGen as the result of cell-line-specific publicly available mutations and fusions.
